# Pan-cancer analysis reveals multifaceted roles of retrotransposon-fusion RNAs

**DOI:** 10.1101/2023.10.16.562422

**Authors:** Boram Lee, Junseok Park, Adam Voshall, Eduardo Maury, Yeeok Kang, Yoen Jeong Kim, Jin-Young Lee, Hye-Ran Shim, Hyo-Ju Kim, Jung-Woo Lee, Min-Hyeok Jung, Si-Cho Kim, Hoang Bao Khanh Chu, Da-Won Kim, Minjeong Kim, Eun-Ji Choi, Ok Kyung Hwang, Ho Won Lee, Kyungsoo Ha, Jung Kyoon Choi, Yongjoon Kim, Yoonjoo Choi, Woong-Yang Park, Eunjung Alice Lee

## Abstract

Transposon-derived transcripts are abundant in RNA sequences, yet their landscape and function, especially for fusion transcripts derived from unannotated or somatically acquired transposons, remains underexplored. Here, we developed a new bioinformatic tool to detect transposon-fusion transcripts in RNA-sequencing data and performed a pan-cancer analysis of 10,257 cancer samples across 34 cancer types as well as 3,088 normal tissue samples. We identified 52,277 cancer-specific fusions with ∼30 events per cancer and hotspot loci within transposons vulnerable to fusion formation. Exonization of intronic transposons was the most prevalent genic fusions, while somatic L1 insertions constituted a small fraction of cancer-specific fusions. Source L1s and HERVs, but not Alus showed decreased DNA methylation in cancer upon fusion formation. Overall cancer-specific L1 fusions were enriched in tumor suppressors while Alu fusions were enriched in oncogenes, including recurrent Alu fusions in *EZH2* predictive of patient survival. We also demonstrated that transposon-derived peptides triggered CD8+ T-cell activation to the extent comparable to EBV viruses. Our findings reveal distinct epigenetic and tumorigenic mechanisms underlying transposon fusions across different families and highlight transposons as novel therapeutic targets and the source of potent neoantigens.

## Main

The role of transposable elements (TEs) in cancer biology has been the subject of extensive investigation across various dimensions. Whole genome sequencing data has greatly facilitated the comprehensive characterization of polymorphic^1–3^ and somatically retrotransposed TEs^4–7^. Some of these retrotransposition events have been implicated as driver events in cancer^6,8,9^. Moreover, a growing body of research is examining chimeric transcripts containing both TEs and genes, referred to here as TE fusions (see Figure S1). Some studies have focused on the phenomenon known as onco-exaptation, where oncogenes initiate transcription via cryptic promoters located within TEs^10–12^. Others have explored exonization events involving TE sequences within transcripts^13,14^. Additionally, there has been a particular emphasis on transcripts containing integrated long terminal repeat (LTR) sequences^15^.

Certain classes of TEs, such as human endogenous retroviruses (HERVs) with their origin in retroviruses^16^, have drawn attention to TEs for their potential role as cancer neoantigens^17^. Peptides derived from TE sequences have been identified in the peptidome presented by major histocompatibility complexes (MHCs) ^18^. Intriguingly, specific HERV expression has been linked to responses to immune checkpoint inhibitors in cancer patients^19^. Most recently, two studies have highlighted the potential of TE fusion transcripts as immunotherapy targets^20,21^. In one study, a pan-cancer analysis focused on chimeric TE transcripts arising from onco-exaptation, revealing that peptides expressed from TE promoters can serve as surface antigens^20^. Another investigation centered on the exonization of TE sequences in lung cancer, demonstrating the resulting peptides recognized by T cells^21^. As previous studies primarily examined limited types of TE fusions derived from TEs annotated in the reference genome, the landscape of various types of TE fusions in human tissues and cancers, especially those originating from polymorphic or somatically retrotransposed TEs remains largely unexplored.

Here, we develop a bioinformatic tool called *rTea* (*RNA-based Transposable Element Analyzer*) to identify TE-fusion transcripts derived from various types of TEs in short-read RNA-seq data. We apply *rTea* to a total of 13,345 uniformly reprocessed RNA-seq profiles, using the same aligner and quality control methods, from four large consortia on Google Cloud Platform (GCP). This enables us to comprehensively map the landscape of TE fusions in human cancers and tissues. Through an integrative analysis, such as with long-read whole genome sequences (WGS) for simultaneous quantification of DNA methylation, we characterize potential epigenetic mechanisms and elucidate tumorigenic roles of cancer-specific TE fusions. Furthermore, we establish a framework to screen neopeptides derived from TE fusions in cancer samples that trigger CD8+ T-cell response and implicate TE fusions as a significant source of potent neoantigens.

### Development of *rTea* to detect transposon-fusion RNA

To investigate the prevalence and functional consequences of transposon-fusion RNA, we developed a computational method, *rTea* to detect TE-fusion transcripts from short-read RNA-seq data. Most methods developed to detect TE-associated events, such as non-reference TE insertions, in genomic data use both clipped reads that span the insertion breakpoints, and discordant reads whose mate reads map to TE sequences at different genomic loci^22^. However, discordant reads supporting TE fusions are rarely present in RNA-seq data that have relatively short read lengths and a much shorter insert size than genome sequencing data (**Figure S2A and B**). Detecting TE fusions from clipped reads alone increases false positive predictions, and the large number of TE-derived reads that map to multiple genomic loci complicates the detection of TE fusions. Thus, we utilized multiple features from aligned reads, such as base quality of clipped sequences, percentage of multi-mapped reads, and matching score of reads to TE sequences to filter out false positives caused by nonspecifically mapped reads (see Methods for details).

We evaluated the detection performance of *rTea* both by using *in silico* simulated data and by experimentally validating predicted fusions using RT-PCR experiments. In the simulation, we created RNA-seq reads containing TE-fusion sequences at different sequencing depths ranging 1X – 200X. As read mapping and fusion detection are more challenging for young TE subfamilies due to high sequence similarity, we used youngest TE subfamilies (*i.e.*, L1HS, AluY, SVA_F, HERV-K, LTR5) in the simulation benchmarking (Details in Methods). *rTea* showed the area under the curve (AUC) for the precision-recall curve of 0.73 – 0.95 for the fusion events simulated at 100X (**Figure S2C, Table ST1**) and showed reliable detection performance with a recall of 0.730, precision of 0.992, and an F1 score of 0.841 when the simulated depth was higher than 10X (**Figure S2D**).

We further evaluated *rTea* by applying it to RNA-seq data we generated from the H1299 non-small cell lung cancer cell line with 108,321,106 reads (TrueSeq RNA v2 100 bp paired-end library and Illumina HiSeq2500 machine from Illumina, CA, USA) and performing RT-PCR validation of predicted TE-fusion events. Among randomly selected 50 events, one failed to amplify, and 34 (69%) produced expected PCR products (**Figure S3, Table ST2**). To examine whether the TE fusions are specific to H1299, we performed RT-PCR for the events validated in H1299 using three additional cancer cell lines (SKBR3, RPMI8226, HCC827) and one normal epithelial cell line (BEAS-2B). We also analyzed in-house and CCLE (Cancer Cell Line Encyclopedia) RNA-seq data^23^ from the three additional cancer cell lines using *rTea*. Six fusions—fusion #12, 21, 34, 42, and 43—called by *rTea* only in H1299 were exclusively confirmed in H1299, absent in the other three cancer and the normal cell lines (**Figure S4A)**. Interestingly, multiple TE-fusion transcripts, including fusion #10 (the exonization of an intronic L1PA5 element in the *CCDC126* gene), were detected in some of the additional cancer cell lines but absent in the normal cell line (**Figure S4A**). We further performed Sanger sequencing of the PCR products of fusion #10 and confirmed the precise fusion breakpoints as predicted by *rTea* (**Figure S4B**). These data suggest the presence of recurrent cancer-specific TE fusions across multiple cancer samples.

### The landscape of TE fusions in human cancers and normal tissues

To understand the extent and type of transposon-derived fusions in human tissues and cancers, we applied *rTea* to RNA-seq profiles generated by four large-genomics consortia: The Cancer Genome Atlas (TCGA) ^24^, the International Cancer Genome Consortium (ICGC) ^25^, the Center for Integrative Omics and Precision Medicine (CoPM) (260 colon cancer samples; unpublished), and the Genotype-Tissue Expression (GTEx) ^26^. The cohorts include 10,257 cancer samples of 34 cancer types and 3,088 normal samples from 28 tissue types. We identified a total of 30,016 TE fusions from normal tissues (mean 203 fusions per sample) and 52,277 TE fusions present in cancer samples but nearly absent in normal tissues (mean 30 fusions per sample), which we call normal and cancer-specific fusions, respectively. Alu elements contribute the most to TE fusions both in normal TE fusions (83%) and cancer-specific fusions (68%), followed by HERV, L1, and SVA elements (**Figure 1A-B**). Compared to normal tissues, the contribution of non-Alu transposons doubled in cancers. For example, L1s and HERVs contributed to 5.7% and 8.4% of normal fusions whereas 12% and 16% of cancer-specific fusions, respectively.

**Figure 1.**
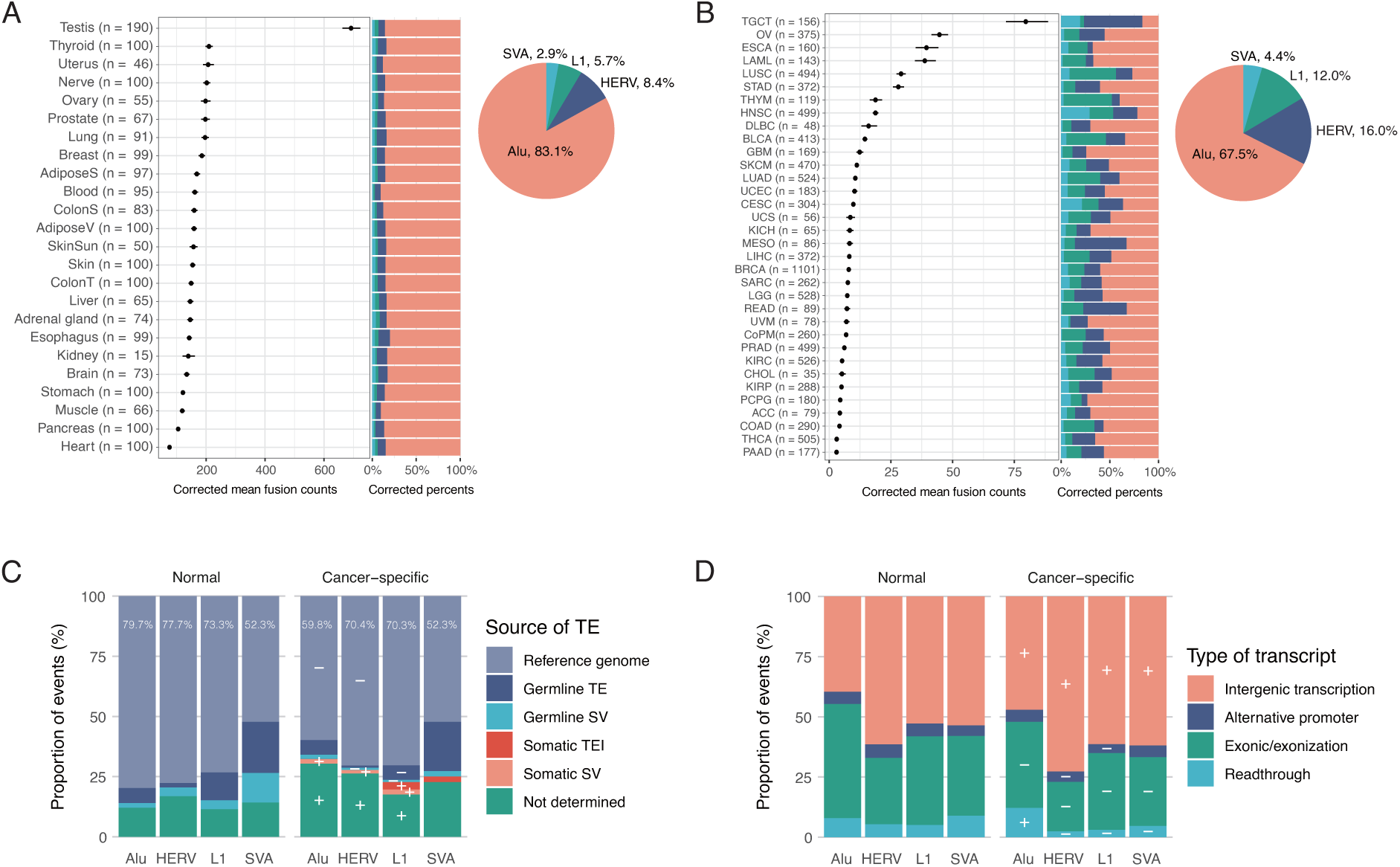
Landscape of TE fusions detected by *rTea*. (A) Corrected mean number of TE fusions per sample detected in GTEx data (2,076 samples). The number of TE fusions were corrected for technical variables, such as read length, and sequencing depth and quality). The error bar represents the 95% confidence interval of the corrected mean value. The pie chart shows the proportion of each TE family. (B) Corrected mean number of cancer-specific TE fusions per sample detected in TCGA (9,645 pan-cancer samples) and CoPM data (260 colorectal cancer samples). Cancer types are labeled using TCGA abbreviations. (C) Types of source TEs in normal and cancer-specific fusions for each TE family. The percentage of reference TEs is labeled for each category. Categories with a significant increase or decrease in cancer-specific fusions compared to normal fusions are marked by ‘+’ or ‘-’, respectively (FDR <0.05, two-sided Fisher’s exact test). (D) Transcript types of normal and cancer-specific fusions for each TE family.

The sensitivity of TE-fusion detection heavily depends on technical differences in RNA-seq data, such as read length, sequencing depth, and sequencing quality, and they are highly variable across and even within cancer and tissue types in the multi-center, long-term large consortia datasets we analyzed (**Figure S5**). In order to estimate the mean TE-fusion count for each tissue and cancer type accurately, we thus used a Generalized Linear Mixed Effect Regression (GLMER) model to correct for the technical variables. Among 28 human tissue types, the testis, where the highest number of genes expressed^27^, showed the highest TE-fusion load (average 639 fusions per sample; **Figure 1A, Figure S6**). Almost half of the TE fusions (48%) found in the testis were not detected in other tissue types, consistent with the distinct epigenetic and transcriptomic signatures of the human testis^28,29^. Interestingly, 77% of cancer samples had at least one testis-specific TE fusions (**Figure S7**). The other tissue types showed a much lower incidence of TE-fusion burden (average 76-210 per sample), with only 5% of them unique to a single tissue type. TE-fusion load exhibited a weak positive correlation with age in most tissues, including visceral adipose, tibial nerve, and esophagus (**Figure S8)**. Among the 24 tissue types, visceral adipose tissue showed the most significant positive correlation between age and the burden of both total TE fusions and Alu fusions (Negative binomial generalized linear model; adjusted p-value = 0.013 and 0.041, respectively), suggesting a potential role of TE-derived fusions in age-related adipose tissue dysfunction^30,31^.

Among 34 caner types, testicular germ cell tumors (TGCT) showed an unusually high burden of cancer-specific TE fusions (mean 80 per sample) (**Figure 1B, Figure S6**). Ovarian cancer (OV), esophageal carcinoma (ESCA), and acute myeloid leukemia (LAML) showed an average of 39-45 fusions per sample, followed by lung squamous cell carcinoma (LUSC) and stomach adenocarcinoma (STAD) with 28-29 fusions on average. Except for AML where no bona-fide somatic L1 retrotransposition has been reported, the cancer types with high TE-fusion loads reportedly showed high rates of somatic L1 retrotransposition^6^.

### Characterization of source TEs creating TE fusions

We characterized the types of source TEs contributing to TE fusions by examining the reads near the fusion breakpoints in WGS data from the same donor as RNA-seq data. First, we created a catalogue of somatic and non-reference germline TE insertions from 1,367 cancer and 1,270 normal tissue samples by collecting a published TE insertion call set from the TCGA and ICGC cohorts^6^, and applying xTea^7^, MELT^32^, and TraFiC^5^ to cancer and matched blood WGS data from 229 CoPM colorectal cancer patients. The catalogue comprises of 15,976 somatic TE insertions (15,900 L1s, 60 Alus, and 16 SVAs) and 2,274,304 non-reference germline TE insertions (1,887,774 Alus, 287,138 L1s, 88,132 SVAs, and 11,260 HERVs). The CoPM colon cancer analysis revealed high rates of somatic TE, predominantly L1, insertions with an average of 36 L1 insertions per tumor; however, the rates were highly variable across individual tumors, consistent with the previous reports^6,33^ (**Figure S9A**). Although most insertions are likely passenger events^6,33,34^, recurrent insertions across multiple patients were found in known cancer genes (**Figure S9B**). Notably, three colon cancer patients (3/229, 1.3%) had exonic somatic L1 insertions in *APC* gene, a well-known tumor suppressor in colon cancer^8,9,35^ (**Figure S9C)**. The two patients had somatic L1 insertions in the last exon (exon 16), producing fusion transcripts detectable in the RNA-seq data. The other patient had a somatic L1 insertion in exon 15, and transcripts from the insertion allele were not detectable in the RNA-seq data, likely due to non-sense-mediated decay.

Most normal and cancer-specific fusions were derived from TEs annotated in the reference genome (78.5% and 63.0%, respectively; **Figure 1C**). A small proportion of TE fusions were derived from non-reference germline TE insertions or other structural variants (SVs). Notably, non-reference SVA insertions accounted for more TE fusions than other TE families both in normal and cancer-specific fusions. Somatic insertions of mainly L1 elements contributed to cancer-specific TE fusions, consistent with somatic L1 mobilization reported in human cancers^4,6^ (**Figure 1C; Table S1**). Overall, TE fusions were derived mainly from reference TE copies, but also from polymorphic and somatic genomic changes involving TEs.

### Distinct splicing hotspots within TEs

TE-fusion events mainly occurred through either the expression of intergenic TEs or the exonization of intronic TEs, with an increased proportion of cancer-specific fusions involving intergenic TEs (**Figure 1D**). Notably, Alu showed more read-through fusions, *i*.*e*., a down-stream Alu being part of the upstream gene transcript, in cancer. We examined the distribution of splice sites within TE consensus sequences and identified distinct hotspot loci across different TE subfamilies (**Figure 2; Table ST3**). The splice donor or acceptor sites were not evenly distributed within TEs but typically confined to specific hotspots. For example, nearly all the splice sites identified in fusions within L1HS elements were splice donor sites near the 5’ end, which a substantial proportion of L1HS copies lack due to frequent 5’ truncation during L1 retrotransposition.

**Figure 2.**
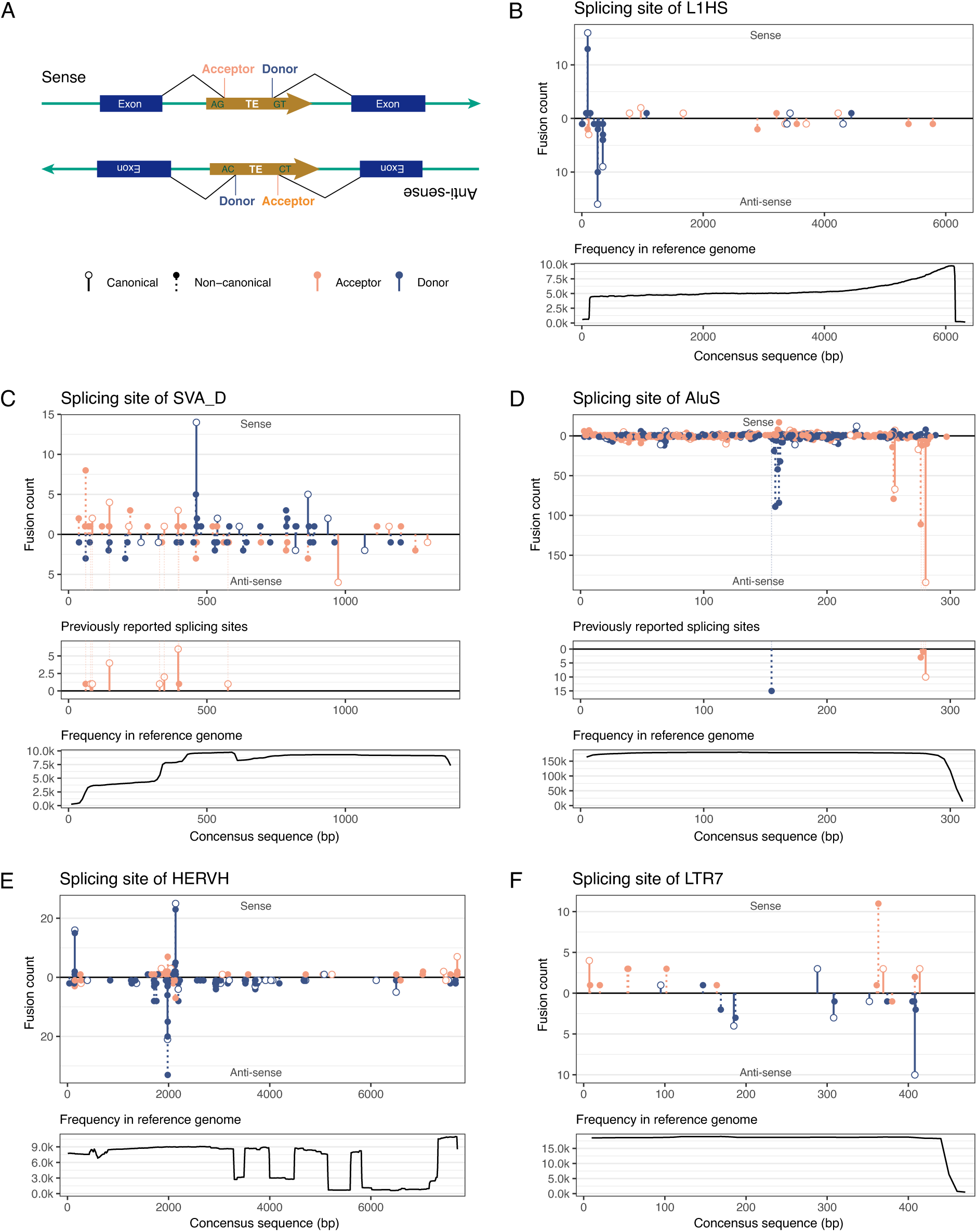
Splicing hotspots within TE consensus sequences. (A) Schematic diagram of an exonized TE with splice acceptor (3’) and donor (5’) sites (orange and blue lollipops, respectively). For TE exonization fusions, the count and type of splice signals are marked within TE consensus sequences for (B) L1HS, (C) SVA_D, (D) AluS, (E) HERVH, and (F) LTR7. Splice sites in the sense and antisense direction are marked upward and downward; canonical (AG/GT) and non-canonical (non AG/GT) splice sequences are marked by solid and dotted line lollipops, respectively. The occurrence of each TE position in the reference genome is shown in the bottom panel. L1HS and SVA_D show expected 5’ truncation patterns. Previously reported splicing hotspots for AluS and SVA_D are shown in the middle panels in C and D.

In contrast, AluS showed both donor and acceptor splice sites on the antisense strand in nearly all reference copies. These splicing hotspots included both canonical (AG/GT) and non-canonical splice signals, although it remains possible that the sequence variation of non-canonical splice sites in the individual samples resulted into canonical splice sequences. Our identification of multiple splicing hotspots aligns with previous findings from the analysis of the human reference genome, expressed sequence tags, and 5’ RACE studies on cell lines^13,14,36^ (**Figure 2).** The presence of splicing hotspots within TEs provide potential therapeutic or intervention targets, for example by antisense oligonucleotide drugs to block cancer-specific TE-fusion formation.

### DNA hypomethylation underlying L1 and HERV fusions in cancer

Previous studies suggested that epigenetic change in TE promoters lead to the expression of oncogenes through a process termed onco-exaptation^10–12^. To determine the relationship between changes in DNA methylation and the formation of diverse types of TE fusions, we compared cancer-specific TE fusions and DNA methylation levels of source TEs from DNA methylation array data generated from 8,365 cancer samples. We found a negative correlation between the number of cancer-specific TE fusions and the mean DNA methylation level in the open sea regions of the genome (areas without CpG islands) for all types of fusions (**Figure 3A**). These negative correlations were not observed in genomic regions containing CpG islands. This association between DNA hypomethylation and increased TE fusions was also confirmed at the individual TE level for all TE families except for Alu (**Figure 3B, Table S2**). Except for Alu, all other cancer-specific TE fusions showed an overall decrease in DNA methylation within 1 Kbp of source TEs (**Figure 3C**).

**Figure 3.**
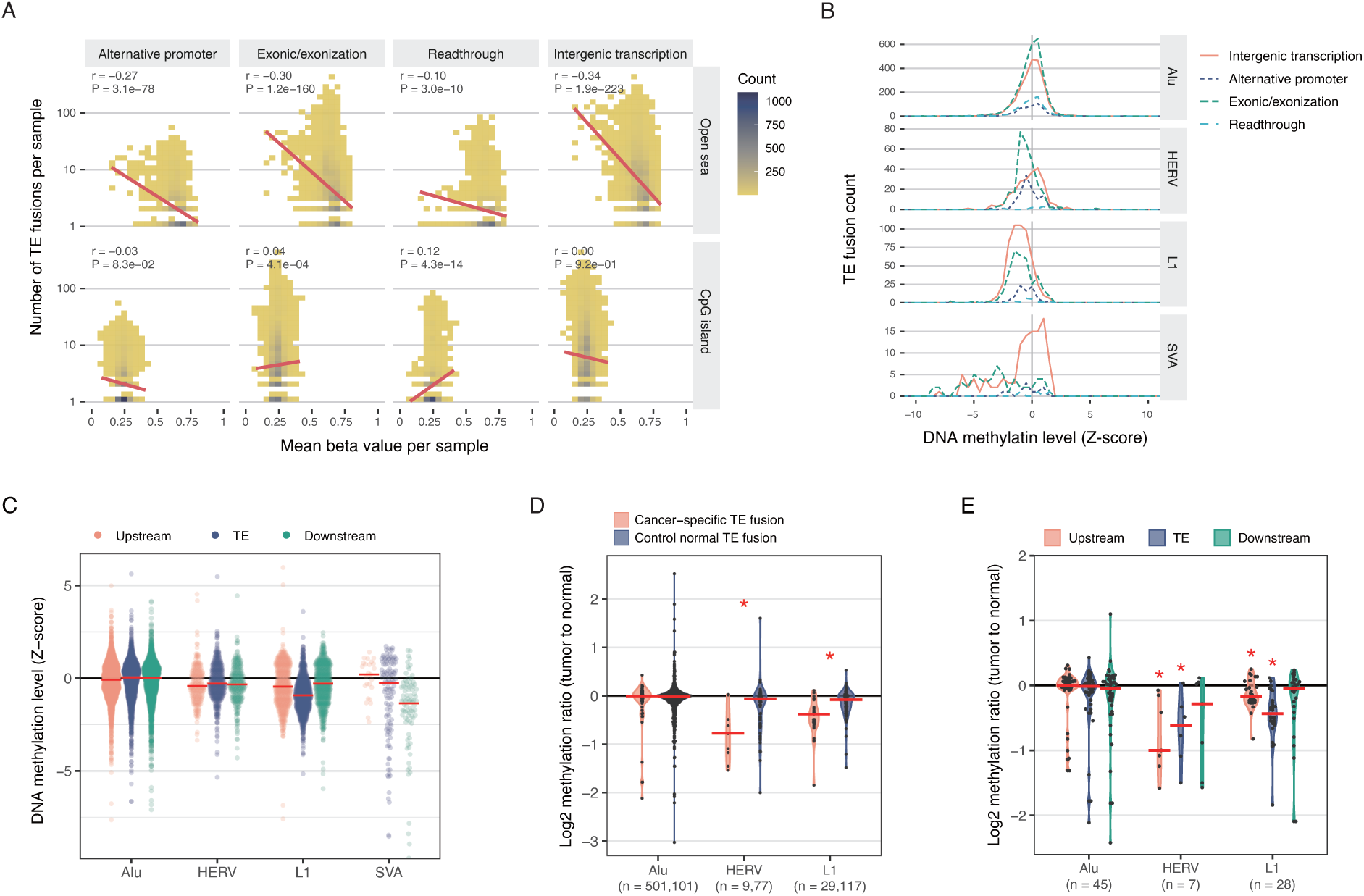
Association between TE fusion and DNA methylation level. (A) Negative correlation between the number of cancer-specific TE fusions and mean DNA methylation level in the open sea. The number of cancer samples per bin is shown as a color scale. The regression line shows the trend. (B) Decreased DNA methylation level near (1 Kbp) source TEs of cancer-specific TE fusions. Z-score of each methylation site was calculated across the same cancer type. Data are shown for TE fusions in TCGA and CoPM samples. (C) TE-family-specific patterns in DNA methylation of source TEs. DNA methylation levels in 1 Kbp upstream, within the TE body, and 1 Kbp downstream of source TEs are marked for cancer-specific fusions detected in TCGA and CoPM samples. (D) Decreased DNA methylation in source HERVs and L1s in cancer fusions observed in ONT long-read WGS data. Tumor to normal log2 methylation ratio was calculated for the source TE involved in cancer-specific (orange) and normal TE fusions (control, blue) detected in five cancer and matched normal pairs with ONT WGS data. Red line represents median; red asterisk indicates a significant difference (P < 0.05, two-sided Wilcoxon rank sum test). N, the numbers of cancer-specific and normal fusions separated by comma in parenthesis. (E) Hypomethylation underlying HERV and L1 fusions in cancer. Tumor to normal log2 methylation ratio was calculated for 1 Kbp upstream, TE body, and 1 Kbp downstream of source TEs for cancer-specific TE fusions from ONT WGS data from five cancer and normal sample pairs. Red line represents median; red asterisk indicates a significant difference (P < 0.05, two-sided Wilcoxon signed-rank test). N, the number of cancer-specific fusions.

To further examine DNA methylation of individual source TEs with higher resolution, we generated long-read WGS data using Oxford Nanopore Technologies (ONT) from five colon cancer and normal tissue pairs. The ONT WGS data allow us to measure DNA methylation levels at individual CpG sites (see Methods). We compared the ratios of DNA methylation levels of the source TEs in the cancer to normal tissue samples and found significantly decreased DNA methylation at source TE loci only for cancer-specific TE fusions involving HERV and L1 (p = 0.014 and < 0.001, respectively; Wilcoxon signed-rank test), but not those involving Alu. The decreased DNA methylation was not observed for TE fusions present in normal tissues (**Figure 3D**). The reduction in DNA methylation was observed in both the upstream and the body of source HERVs and L1s (**Figure 3E, Table S3**), but not in the downstream of any TEs. No reduction in DNA methylation near or within Alus was confirmed in this ONT WGS-based DNA methylation analysis as well. Alu is unique among TEs in that it is transcribed by RNA polymerase III^37^, which might be less affected by DNA methylation. Taken together, these results suggest that DNA hypomethylation upstream of or within the TE body lead to TE-derived fusions, except for Alus, in cancer.

### Contribution of TE fusions in tumorigenesis and prognosis

To understand the role of TE fusions in tumorigenesis, we examined the prevalence of cancer-specific fusions in known cancer genes. Overall, cancer-specific TE fusions were enriched in cancer genes relative to normal TE fusions. Specifically, out of 16,492 cancer-specific TE fusions found in protein coding genes, 845 (4.9%) involved known cancer genes, while out of 11,446 normal TE fusions in protein coding genes, 449 (3.8%) involved cancer genes (p < 0.001; Logistic regression). Interestingly Alu fusions were enriched in oncogenes suggesting they presumably lead to gain of function or overexpression of oncogenes; L1 fusions were enriched in tumor suppressor genes, but slightly depleted in oncogenes suggesting L1 fusions are likely to contribute to tumorigenesis by loss of function of tumor suppressor genes (**Figure 4A**).

**Figure 4.**
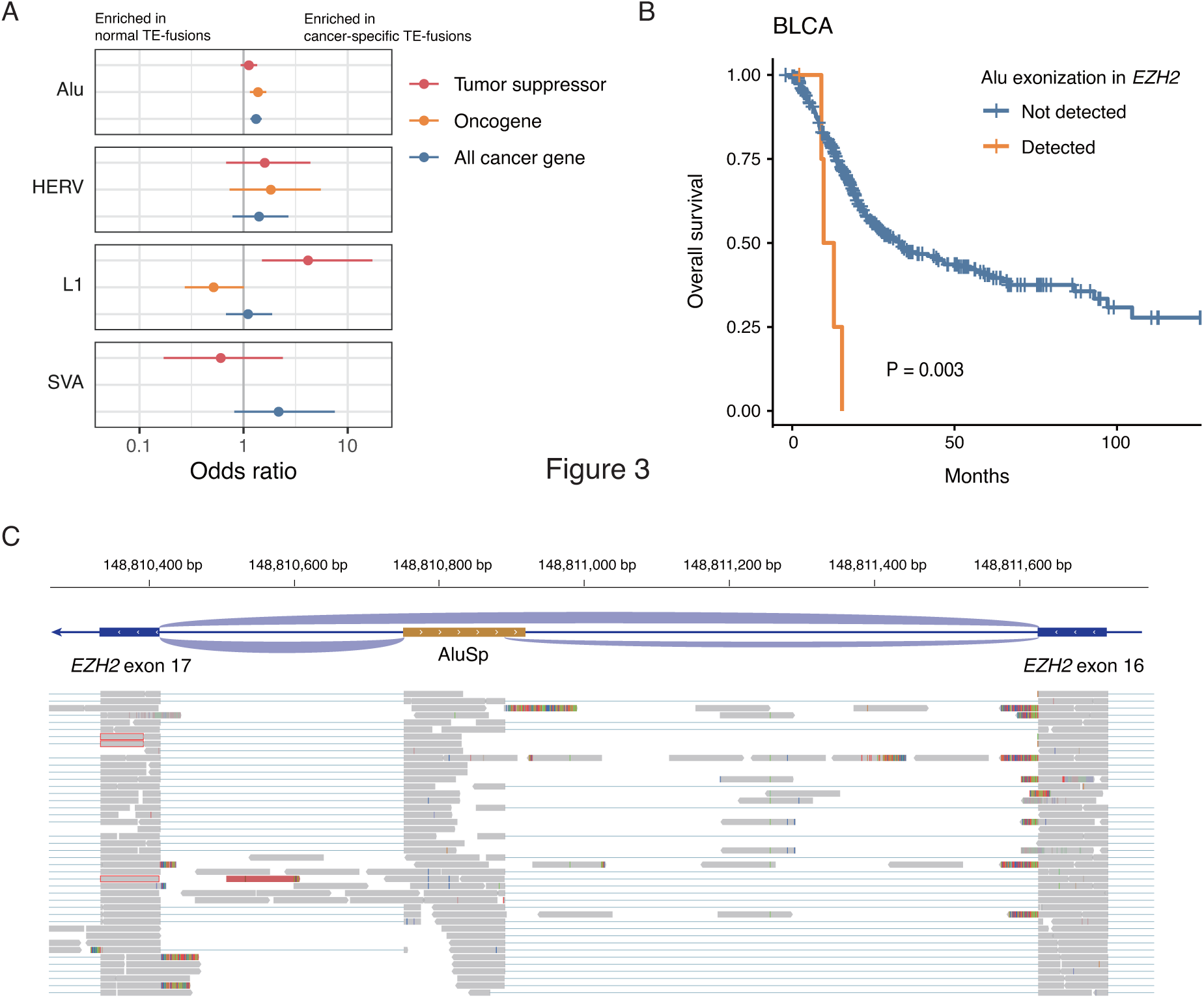
TE fusions enriched in known cancer genes and associated with patient survival. (A) Odds ratios representing tumor suppressor genes (red), oncogenes (orange), and all cancer genomes (blue) are marked for cancer-specific TE fusions for each TE class. (B) Survival rates of patients with and without the AluSp exonized *EZH2* gene in the TCGA bladder cancer cohort. (C) Schematic and Integrative Genomics Viewer (IGV) screenshot of the exonization of an intronic AluSp in *EZH2* gene detected by *rTea*. Reads with split mappings (blue lines) show both splice junctions—exon 16 and AluSp as well as AluSp and exon 17. A pileup of reads (gray box) on the AluSp shows exonization of the intronic TE.

We also examined whether cancer-specific fusion loads correlate with patient survival and found varied patterns across TE family and cancer type. Cancer-specific fusion loads were more often associated with poor prognosis (*e.g.,* the number of Alu fusions in ovarian serous cystadenocarcinoma) than better prognosis (**Figure S10**). We found a recurrent, cancer-specific fusion between L1PA2 and the proto-oncogene *MET* in 93 cancer patients of diverse cancer types (Fusion ID: dup4642; **Figure S11**), which has been reported to be associated with poor prognosis in various cancer types^38–40^. We also found that a cancer-specific exonization event in the *EZH2* gene from a reference AluSp within intron 16 was observed in multiple bladder cancer patients, and patients with the fusion showed poor survival (Hazard ratio [95% confidence interval], 4.3 [1.6 – 11.8]; **Figure 4B and C**). EZH2 is a histone methyltransferase involved in regulating cell division and known to contribute to cancer development through gain of function mutations or overexpression,

### TE fusion activating T-cell immunity and shaping immune microenvironment

TE fusions often produce novel peptide sequences that have the potential to trigger T-cell immunity^15^. To investigate whether TE fusion-derived peptides can be processed as neoantigens and invoke T-cell response, we used MuPeXI^41^ to identify peptides from TE fusions identified in the CoPM dataset capable of bind to major histocompatibility complex class I (MHC-I) and dissimilar enough to normal human peptides to induce an immune response. To reduce false positive predictions, we additionally employed a structure-based peptide-MHC-I binding prediction method to remove peptides unlikely to bind and any peptides where no mutated residue is accessible to bind T cells (see Methods). We selected 33 peptides predicted to bind to five HLA alleles (HLA-A*02:01, A*24:02, B*58:01, C*14:02 and C*54:01) based on the population frequency of HLA class I and TE fusion types. Of these, 20 TE-derived peptides showed detectable binding signals in the *in vitro* MHC-I binding assay (**Figure 5A**). The binding experiments showed that the combination of MuPeXI and structure-based filtering is highly predictive of actual binding with more strict thresholds (NetMHCpan percentile from MuPeXI < 0.5 and binding energy Z-score < 0) (**Figure 5B**).

**Figure 5.**
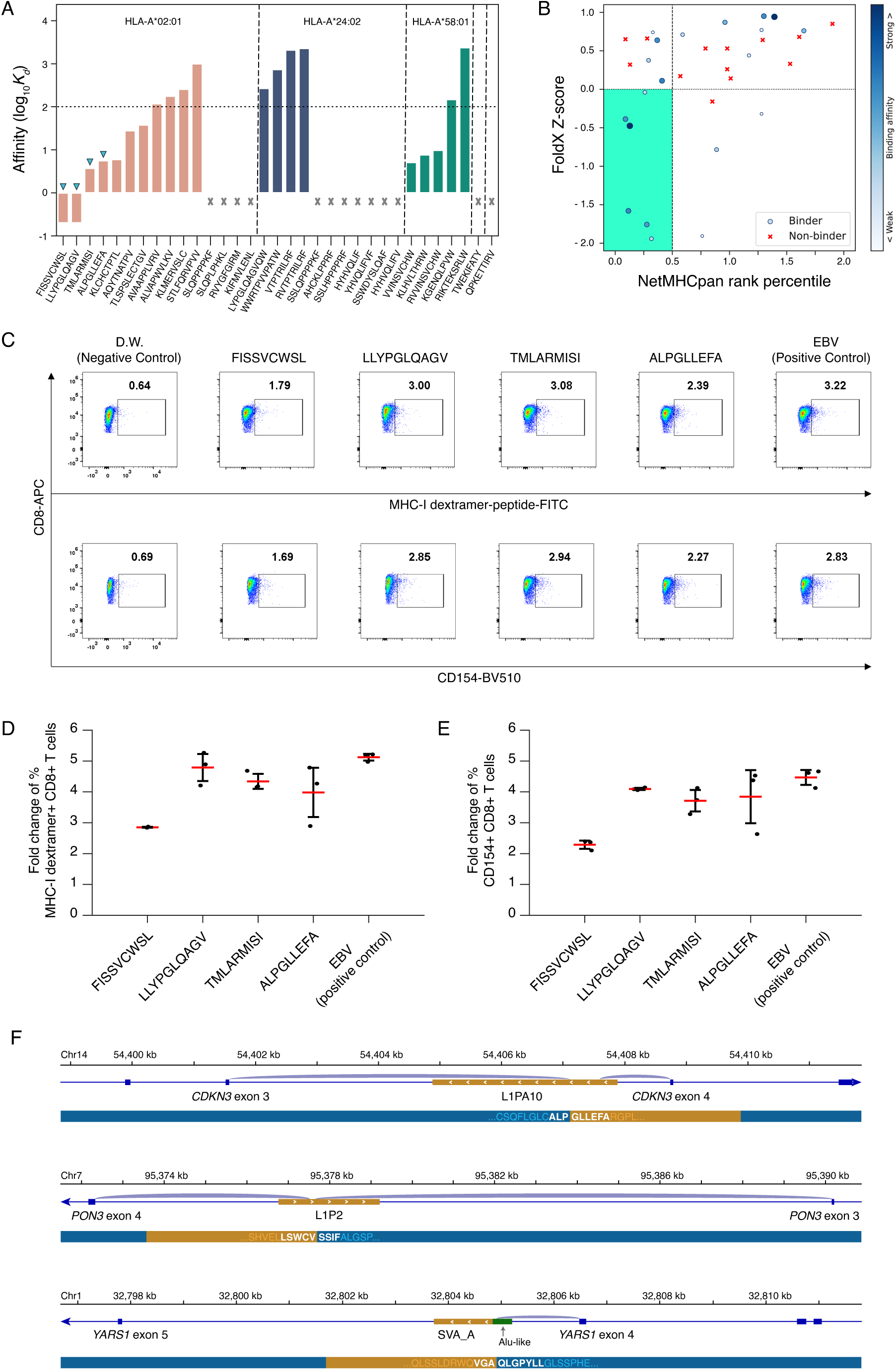
Immunogenicity of TE-derived peptides. (A) Binding affinity of TE-derived peptides to major histocompatibility complex class I (MHC-I) molecule. TE-derived peptides predicted to bind to MHC-I were selected from CoPM colorectal cancer samples using *in silico* affinity prediction and structural modeling. Among the 33 peptide candidates, 20 peptides exhibited binding signals. The four peptides with the highest binding affinities to HLA-A*02:01 were further tested for T-cell activation. (B) Prediction power of computational methods to predict MHC-I binding of TE-derived peptides. NetMHCpan rank percentile is a prediction score for a peptide to be presented on MHC-I, normalized by comparing the score to the prediction of random peptides. FoldX Z-score represents the Z-score of predicted binding affinity calculated from structural modeling. We found that the structural filtering alone can be extremely predictive if Z-score < 0. The combination with NetMHCpan (rank percentile < 0.5) may further remove weak binders. (C) Detection of CD8+ T cells that specifically bind to HLA-A*02:01-presented peptides. Healthy donor CD8+ T-cells specifically binding to HLA-A*02:01-presented peptides were cultured for 21 days during co-culture with autologous peripheral blood mononuclear cells (PBMCs) and peptide-pulsed T-cells. Peptide-specific T-cells were detected with fluorescein isothiocyanate (FITC)-labeled MHC-I dextramer. The activation of T-cells was assessed by measuring CD154 expression. (D) Quantification of peptide-specific CD8+ T cells which bind peptide-MHC-I dextramer. The fold change was calculated by comparing the percentage of positive cells with distilled water-treated negative control. (E) Quantification of CD8+ and CD154+ expression on healthy donor T-cells. The fold change was calculated by comparing the percentage of positive cells with distilled water-treated negative control. (F) Configuration of TE fusions from which T-cell recognized peptides were derived. The peptides ALPGLLEFA, FISSVCWSL, and LLYPGLQAGV can be produced from the TE fusion of CDKN3-L1PA10, PON3-L1P2, and YARS1-SVA_A, respectively. The red line and error bar in (D) and (E) represent mean and standard deviation, respectively

To confirm that these peptides are targeted by T cells, we selected the four strongest binders to HLA-A*02:01 to perform T-cell binding assays using human peripheral blood mononuclear cells (PBMCs). We pulsed these four peptides into autologous PMBCs co-cultured with healthy donor-derived CD8+ T cells that specifically bind to peptides presented by HLA-A*02:01 and allowed the T cells to proliferate for 21 days. We measured the activation and proliferation of T cells both using peptide-specific staining by fluorescein isothiocyanate (FITC)-labeled peptide dextramer staining, and using the general T-cell activation marker CD154 (CD40 ligand). All four peptides led to robust activation and proliferation of T cells (**Figure 5C-E**). All but one peptide led to activation and proliferation comparable to a positive control peptide generated from the Epstein-Barr virus, suggesting endogenous transposon-derived peptides can be as immunogenic as foreign or viral peptides.

We examined where exactly the immunogenic 9-mers originated within the fusion transcripts. Two of these peptides, FISSVCWSL and ALPGLLEFA, were derived from the junctions between the exon 3 of *PON3* and *CDKN3* genes, and the exonized reference L1PA10 and L1P2 within the introns of *PON3 and CDKN3*, respectively (**Figure 5F**). The LLYPGLQAGV peptide was generated from the junction between exon 4 and the exonized SVA_B in the intron of the *YARS1* gene. The TMLARMISI peptide originated from the antisense internal sequence of AluSc. In our analysis, this peptide was produced from the fusion with the *MRPL3* gene. Since, there are 35,544 AluSc copies in the reference human genome. Expression of the antisense sequence from any of these copies through the formation of TE fusions can result in the generation of this peptide.

Beyond our proof-of-concept experiments for antigenic potential of TE-derived peptides, several studies including our previous work suggested that reactivated transposons trigger innate immune responses^17,18,20,42^ and might play a more complex, long-term immune-modulatory role in primary tumors leading to the observation of anti-correlation between cancer immunity and somatic retrotransposition rate^33^. Here, we examined the relationship between cancer-specific TE fusions and the immune microenvironment by measuring the correlation between the number of cancer-specific TE fusions and the tumor immune score calculated using the ssGSEA^43^ method with marker gene sets^44^. Across numerous cancer types, we found a significant negative correlation between the number of Alu or L1 fusions with nearly every immune component, with less widespread negative correlations for HERV and SVA fusions (**Figure S12**). The only notable exception is the positive correlation observed between the number of SVA fusions and immune score in testicular germ cell tumors. The testis is normally immune privileged, and the positive correlation may indicate immune infiltration within the testicular tumor.

## Discussion

Our pan-cancer analysis highlights the prevalence of TE-fusion transcripts in both cancer and normal tissues along with the potential roles they play in tumorigenesis and immunogenicity. While the overall number and proportion of the TE families involved varied, we detected cancer-specific TE fusions in every cancer type. Except for the Alu family of TEs, which are under the control of RNA polymerase III, we found that cancer-specific TE fusions are associated with DNA hypomethylation upstream of or within the TE bodies in cancer, suggesting epigenetic aberration underlying some TE-fusion formation. Additionally, we found these cancer-specific TE fusions are enriched in known cancer genes, with Alu and L1 family TEs showing enrichment in oncogenes and tumor suppressor genes, respectively, which can drive tumor development and alter prognosis by distinct dysregulation mechanisms from one another. For example, we found a recurrent exonization of an intronic AluSp element between exons 16 and 17 of the *EZH2* gene that leads to worse prognosis for patients with bladder urothelial carcinoma.

We also found a negative correlation between the number of TE fusions and the intensity of the immune response in the tumor microenvironment (measured by the immune score) in most cancer types, *i.e.*, the fusion rate is highest in tumors with the least immune activity. This trend is also present in normal tissue where we found the testis, an immune privileged tissue, had ∼3 times more TE fusions than any non-immune privileged tissues. This observation is consistent with previous work showing that epigenetic cancer therapies, including the induction of DNA hypomethylation, lead to increased TE expression and activation of the innate immune system^45–47^. However, the expression of normally suppressed TEs can also act as a source of neoantigens for the adaptive immune system and trigger immune infiltration into tumors, leading to better outcomes for immunotherapies^5,18,48^. In addition, despite the absence of direct experimental evidence, it is reasonable to hypothesize that an elevated TE-fusion level could potentially lead to a compromised immune activity. More investigation is warranted to understand whether an increase in TE-fusion transcripts play an active immunomodulatory role in human tissues and diseases.

The novel peptide sequences produced by TE fusions provide an additional source of neoantigens that were previously predicted to bind to MHC-I^6,15^ and lead to T-cell activation^48^. Here, using fusions detected in colon cancer samples from the CoPM consortium, we demonstrated that computationally predicted peptides derived from the junction sites of TE fusions as well as peptides internal to the TE sequence within the fusion transcripts can bind to MHC-I. Further, we demonstrated that these peptides stimulate T-cell proliferation, generating responses comparable to the Epstein-Barr virus-derived immunogenic peptide. This confirms their potential role as neoantigens for immunotherapy. While we identified a large diversity of unique TE fusions across cancer types, splicing hotspots within the TE sequences and the immunogenicity of internal TE sequences raise the possibility that multiple unique TE-fusion events, potentially created by heterogeneous epigenetic and genomic cues, generate the same antigenic peptides. These splicing hotspots can also facilitate the development of antisense oligonucleotide therapies to block splicing formation in oncogenes or tumor suppressor genes in conjunction with traditional treatments.

The prevalence of TE fusions in normal tissues also suggests that they play a role in normal cellular functions as well, as has been described for an alternative promotor fusion driving isoform expression that controls the timing of pre-implantation development in mammals^49^. However, the mis-regulation of genes through the expression of TE fusions, as seen in the enrichment of TE fusions in tumor suppressor genes and oncogenes, can potentially disrupt other cellular functions leading to other disease states. Additionally, the immunogenicity of the novel peptides produced from these transcripts raises the possibility of their involvement in inflammatory and autoimmune diseases. Further research is needed to determine the full scope of the role TE fusions play in human health and disease.

## Methods

### RNA-based Transposable Element Analyzer (rTea)

#### TE fusion detection

We developed *rTea* (*RNA-based Transposable Element Analyzer*) to detect TE fusions in RNA sequencing data (https://github.com/ealeelab/rtea). It takes a BAM alignment file of a FASTQ file as input. It converts the BAM file to FASTQ files using Picard (ver. 2.21.7) and trims the adapter sequences using fastp (ver. 0.20.0) ^50^. Then, it aligns the trimmed reads to the reference genome version GRCh38 using HISAT2 (ver. 2.1.0) ^51^. To allow multiple mappings and prevent the generation of very large gaps, the parameters for HISAT2 were customized (--sp 1,0 --score-min L,0,-0.5 --pen-canintronlen S,9,0.1 --pen-noncanitronlen S,9,0.1 --max-intronlen 90000 --dta -k 10).

The mapped BAM file was sorted and indexed using Samtools (ver. 1.9) ^52^. The clipped reads were collected using the Bamtools API (ver. 2.5.1) and mapped to the reference TE sequences using BWA (ver. 0.7.17) ^53^. The reference TE sequences contain Repbase^54^ or Repeatbrowser^55^ consensus sequences (Supplementary Document 1: consensus sequences per each TE type) and the reference TE sequences annotated by RepeatMasker (http://www.repeatmasker.org) with the following criteria: All AluYs with size >300 bp and divergence < 5%; all L1Hs with size > 6kb and divergence < 5%; all SVA (SVA_A, SVA_B, SVA_C, SVA_D, SVA_E, and SVA_F) with size > 2kb and divergence < 10%; and all HERV (HERVK-full, HERVK, HERVK11, HERVK11D, HERVK13, HERVK14, HERVK14C, HERVK22, HERVK3, HERVK9, HERVKC4, LTR3, LTR3A, LTR3B, LTR5, LTR5_Hs, LTR5A, LTR5B, LTR13, LTR13A, LTR14, LTR14A, LTR14B, LTR14C, LTR22, LTR22A, LTR22B, LTR22B1, LTR22B2, LTR22C, LTR22C0, LTR22C2, LTR22E, MER9a1, MER9a2, MER9a3, MER9B, MER11A, MER11B, MER11C, MER11D, HERVH-full, HERVH, HERVH48, LTR7, LTR7B, LTR7C, LTR7Y, HERV17, LTR17, HERVL, HERVL18, HERVL32, HERVL40, HERVL66, HERVL74, MLT2A1, MLT2A2, MLT2B1, MLT2B2, MLT2B3, MLT2B4, MLT2B5, MLT2C1, MLT2C2, MLT2D, MLT2E, and MLT2F) with size > 300 and divergence < 5%.

*rTea* uses the Bioconductor package (ver. 3.10) of R software (ver. 3.6.2) to analyze the clipped read alignments. Genomic positions with > 3 clipped reads where clipped sequences mapped to reference TE sequences were selected as candidate TE fusion positions. The longest clipped sequence at each candidate position was kept as the representative clipped sequence. Candidate positions were excluded if the clipped sequence was a simple repeat or from a poly-A tail, or did not align to the consensus TE sequence. With the alignment options described above, HISAT2 tends to clip reads containing a single nucleotide polymorphism. These inappropriately clipped positions were also excluded.

For the remaining candidate positions, we collected clipped reads that map to the TE sequences and extracted several features to examine the confidence of the fusion transcript. We considered a position low confidence if any of the following criteria were met:

- Unique supporting reads < 3
- Unmatched clipped reads ≥ 10 × matched clipped reads
- Median base quality of clipped sequence ≤ 25
- Percent of reads clipped on both sides ≥ 40%
- Percent of multi-mapped reads = 100%
- Percent of low mapping quality reads = 100%
- Average alignment scores of reads that map to consensus TE sequences ≥ 0

All positions that do not meet any of these criteria are considered high confidence for TE fusions. If the clipped position occurs in a repeat region, a substantial number of reads can be aligned to that position. To save memory and time, reads were down sampled to a maximum of 100,000 when collecting the clipped reads for each position.

To annotate each fusion event, clipped reads containing matched TE sequences were collected and compared with the reference transcriptome using R package EnsDb.Hsapiens.v86 (ver. 2.99.0). The intronic and exonic gaps were matched and the transcript with the highest matched read count was selected as the chimeric transcript. The TE fusions were annotated as follows:

- ‘Alternative transcription start’ – The TE-gene junction was located upstream of the matched transcript
- ‘Exonic/exonization’ – The TE-gene junction was located within the genomic position of the matched transcript
- ‘Readthrough transcription’ – The TE-gene junction was located downstream of the matched transcript
- ‘Intergenic transcription’ – No matched reference transcript identified

Clipped sequences were additionally aligned with the nearby reference genomic sequence to determine whether the TE sequence originated from a TE present in the reference genome. If the mate-read from the paired-end sequencing was mapped to a nearby (within 100,000 bp) genomic position, TE sequences were aligned with the reference genomic sequence up to the position of the mate-read. Otherwise, reference genomic sequences up to 100,000 bp from the clipped read were used for alignment.

#### In silico validation of rTea using simulation RNA-seq

The youngest subfamilies of the target TE families (L1HS, AluY, SVA_F, HERV-K, LTR5) were selected for validation. For each TE subfamily, 1,000 TE sequences were sampled from RepeatMasker annotated genomic sequences with length > 100 bp and percent of deviation < 5%. Then these TE sequences were inserted into a random position of a transcript randomly selected from the reference transcriptome (Ensembl release-94). Five sets of 100 exonic and 100 intronic TE insertions were generated. Background FASTQ reads without TE fusions were generated based on the reference transcriptome sequences using wgsim (https://github.com/lh3/wgsim) based on the reference transcriptome sequences containing 80-90 million reads. The read counts of background expression were based on 5 GTEx samples downloaded from the GTEx Portal (https://gtexportal.org/home/datasets): blood (GTEX-OXRL-0005-SM-3LK6A), esophagus (GTEX-13X6K-2126-SM-5O9D4), colon (GTEX-Z93S-2626-SM-57WBX), lung (GTEX-1GMR8-1426-SM-7RHHL), and brain (GTEX-145MH-3026-SM-5Q5DZ). For the TE inserted transcripts, we generated reads to a depth of 1, 2, 3, 4, 5, 7, 10, 20, 30, 50, 100, and 200X and combined with background FASTQ reads. These simulation data were made for two conditions: read length of 100 bp, insert size of 167 ± 20 bp (comparable sequencing condition to the CoPM and the ICGC data), and read length of 75 bp and insert size of 119 ± 10 bp (comparable sequencing condition to the majority TCGA data). Using the simulated FASTQ files, TE fusions were detected by *rTea*. The *rTea* results were classified as true positives if both the breakpoint position and TE-fusion sequence matched with the generated insertion breakpoint and TE sequence. The two sides of TE-fusion junctions were counted separately. Precision, recall, and F1 were calculated to measure the performance. With 100X data, the precision-recall curve was drawn by changing the cutoff value for the supporting read counts and the area under the curve (AUC) was calculated.

#### Experimental validation of rTea with RT-PCR assays

We generated RNA-seq data from the H1299 non-small cell lung cancer cell line with 108,321,106 reads (TrueSeq RNA v2 100 bp paired-end library and Illumina HiSeq2500 machine from Illumina, CA, USA). *rTea* was run on the RNA-seq data, and 50 TE fusions were randomly selected for experimental validation (**Table ST2**). TE-fusion-supporting reads were assembled by CAP3^56^ to know the exact sequences around the junction of TE fusions. RT-PCR primers were designed to detect the TE-fusion sequences. The TE fusions detected by RT-PCR were further tested using four additional cell lines (SKBR3, breast cancer; RPMI8226, HCC827, multiple myeloma; and BEAS-2B, normal bronchial epithelium). RNA-sequencing and *rTea* analysis was done on the SKBR3 and HCC827 cell lines. CCLE RNA-sequencing data was also downloaded for SKBR3, RPMI8226, and HCC827 cell lines used as input for *rTea*.

The five cell lines, H1299, SKBR3, RPMI8226, HCC827, and BEAS-2B, were obtained from ATCC (Manassas, VA, USA). For each region of interest, TE-fusion sequences were amplified by PCR using the primers listed in **Table ST2**. Total RNA was isolated from the cells using the RNeasy mini kit (Qiagen, Hilden, Germany) and used to synthesize cDNA with the SuperScript III First-Strand Synthesis System (Life Technologies, Carlsbad, CA, USA) and 20-mer oligo(dT) primers. cDNA was amplified by PCR using the primers with Herculase II Fusion DNA polymerase (Agilent Technologies, Santa Clara, CA, USA). PCR products were separated by electrophoresis through a 2% agarose gel in 1× TBE and stained with GelRed (Biotium, Hayward, CA, USA). To confirm the sequence of each band, the PCR products were gel purified using the QIAquick Gel Extraction kit (Qiagen, Hilden, Germany) and verified by Sanger sequencing.

### TE-fusion detection using large-scale consortium data

#### RNA-seq, short- and long-read WGS, and DNA methylation datasets

We collected RNA-seq data from 3,088 normal sample and 10,257 cancer samples, WGS data from 1,367 tumor and matched normal samples, and DNA methylation array data from 538 normal sample and 8,365 cancer samples from four consortium projects. The projects are: Genotype-Tissue Expression (GTEx), the Cancer Genome Atlas (TCGA), the International Cancer Genome Consortium (ICGC), and the Center for Integrative Omics and Precision Medicine (CoPM). From GTEx, we analyzed RNA-seq datasets of 2,076 normal tissue samples from 28 tissue types. From TCGA, we analyzed a total of 10,356 RNA-seq datasets (9,645 cancer and 711 normal samples), 786 WGS datasets, and 8,349 DNA methylation arrays (7,966 cancer and 383 normal) for 33 cancer types. From the ICGC, we downloaded 434 RNA-seq (352 cancer and 82 normal), 352 WGS, and 186 DNA methylation arrays datasets of five cancer types.

The CoPM is a cancer genomics project on colon cancer in South Korea among Samsung Medical Center, Seoul National University Hospital, Asan Medical Center, Seoul St. Mary’s Hospital, and Seoul National University Bundang Hospital (http://eng.pmi.re.kr/g5/). From the CoPM, we used bulk RNA-seq data for 260 cancer and 219 matched normal tissue samples, WGS data for 229 cancer and 229 matched normal tissue samples, and DNA methylation array data for 213 cancer samples and 155 normal samples from CoPM. In addition, we generated long-read WGS data for 5 pairs of tumor and matched normal tissue samples using Oxford Nanopore Technologies (ONT). The ONT WGS data allows us to quantify genome-wide DNA methylation level (see DNA methylation analysis for details). The raw RNA-seq and WGS data from the CoPM cohort have been deposited in the KoNA (https://kobic.re.kr/kona). The data information is provided in the data availability section.

#### GenomeFlow for parallel genomic data processing on GCP

In order to analyze a large number of RNA-seq data from multiple consortia, we developed GenomeFlow (a manuscript in preparation), a job and task optimization and scheduling tool on Google Cloud Platform (GCP). We utilized Google Kubernetes Engine (GKE) as the computational architecture for running *rTea* in distributional processing. GenomeFlow was written as a Python (ver. 3.6) script using Google API Core (ver. 1.17.0). GenomeFlow deploys a task scheduler service and controller based on RabbitMQ (ver. 3.7) and creates a Cloud SQL database for a MySQL instance (ver .5.7) to manage jobs and tasks. The database is used to record successes, failures, and pending tasks to prevent duplication. GenomeFlow prepares each task, injects into the user input docker file to control the scheduler and management database, and establishes cloud storage to store the results. Each task uses the asynchronous blocking connection of Pika (ver. 1.1.0). A task involves downloading a sample, running the user input commands, storing the result file from the command, and recording the processing status. Once the workflow file (written in WDL/Snakemake syntax) is submitted by a user, GenomeFlow calculates the optimal parameters for CPU, memory, and disk allocation. GenomeFlow then runs the job for all samples specified in the input configuration.

#### Combining and filtering the rTea results

To aggregate TE fusions from individual samples, we performed a pairwise comparison of TE fusions in the *rTea* results from each sample. Fusions with breakpoints within a 5bp range and 70% sequence identity in their clipped regions were combined into a set of non-redundant fusion events. Events where ≥ 50% of the candidate TE fusions are high confidence were kept in the analyses. TE fusions were classified as cancer-specific when they were found exclusively in cancer samples and absent in all normal tissue samples, or if they were present in < 0.1% of all normal tissue samples with a significantly higher frequency in tumor samples using the Chi-squared test.

### Pan-cancer TE-fusion analysis

#### Estimation of TE-fusion count per cancer or tissue type adjusted by technical variables

We considered five technical variables of RNA-seq library preparation and sequencing that can affect TE-fusion detection: Q30 rates, median insert size, mean read length, number of total reads, and duplication rate. These technical variables differ broadly across cancer and tissue types as well as individual sample level (**Figure S5**). To make the number of detected fusions from each sample comparable, we calculated these technical variables to correct the estimated mean number of TE fusions per sample. We calculated the mean numbers of TE fusions for normal tissue samples from the GTEx data and the mean numbers of cancer-specific TE fusions for tumor tissue samples using the TCGA and CoPM data. A synthetic variable, mean read length × (1 - duplication rate) × number of total reads, was also considered. We also tested the importance of each feature to model the observed number of TE fusions using a Generalized Linear Model (GLM).

We used a Generalized Linear Mixed Effect Regression (GLMER) model to correct the mean TE-fusion count for these technical variables. We used the rstanarm (ver. 2.21.1) package in R (ver. 4.0.2) to make the GLMER model. We used the covariates as fixed effects, the sample group (tissue and cancer type) as random effects with the formula TE-fusion count ∼ covariates

+ 1|sample group. We trained the model using a Bayesian approach.

#### Association analysis with TE-fusion count

We analyzed the association between TE-fusion counts and age. We built a negative binomial generalized linear model (nb-GLM) with the lme4 package (ver. 1.1-26) in R (ver. 4.0.2). We made a total of 120 models for 24 tissue types and Alu, L1, HERV, SVA, total fusion counts. We measured the significance of the association between TE-fusion count and age with the following formula: TE-fusion count ∼ covariates + age. P-values obtained from the models were adjusted to produce FDRs.

We calculated the number of expressed genes based on the transcripts per million (TPM) values in the GTEx Analysis V7. The TPM values were converted to 0 (not expressed) for TPM values < 1 and to 1 (expressed) for TPM values ≥ 1. For each TE family, we calculated the Pearson correlation with p-values between the number of TE fusions and the number of expressed gene in the same sample. We converted FPKM values for genes in TCGA, ICGC, and CoPM datasets to TPM for consistent analysis.

#### Splicing hotspot analysis

We selected exonization-type TE fusions with an identifiable source TE for splice site analysis. The subfamily of a source TE was identified by matching the position of the source TE with the RepeatMasker-annotated reference genome. The sequence at the splice site for the TE region was aligned to the consensus TE subfamily sequence downloaded from the UCSC Repeat Browser^55^. Splice site positions along the consensus TE sequence were collected for each TE subfamily. The donor or acceptor site for each splicing event was determined by the position and strand of TEs relative to gene. We compared splice hotspot loci we identified with previously reported hotspots^13,14,36^.

### Association analysis with DNA methylation

#### DNA Methylation array data

We downloaded probe information for the DNA methylation array from the Illumina Support Center (https://support.illumina.com/). For each sample, the mean beta values were calculated for all regions, CpG sites, and open sea regions. The Spearman rank correlation between the corrected number of cancer-specific TE fusions per sample and the mean beta value was calculated for each type of TE fusions. To determine the effect of DNA methylation on individual TE fusions, probes located within 1000 bp of the source TE were selected, and the Z-score was calculated among the beta values of the same probes in samples of the same cancer type. For each TE class and TE fusion type, the two-sided Wilcoxon signed-rank test was used to test whether the Z-score significantly deviated from 0.

#### Methylation calling for ONT WGS data

For preprocessing, we generated raw FASTQ files with five FASTA files from Oxford Nanopore Technologies (ONT) using the nanopolish (ver. 0.9.2) indexing option. Then, we checked the FASTQ files and removed any duplicate reads using a custom python script. We used minimap2 (ver. 2.13-r852) to align the base called reads to the hg38 reference genome with the -x map-ont option. We then used nanopolish index option to make index files of FASTQ files. We then used calculate_methylation_frequency.py scripts of nanopolish to calculate the methylation frequency (0-1) of CpG sites.

#### Analysis of DNA methylation changes near TE fusions using ONT WGS data

We measured the tumor to normal DNA methylation ratio from ONT WGS data by dividing the methylation level from a tumor sample by the methylation level from the matched normal tissue sample. The 1,000 bp flanking region of a source TE was defined to observe the methylation pattern of the upstream and downstream regions. We calculated the methylation level of each region using the CpG methylation obtained through nanopolish. We measured the average methylation rate in consideration of the number of CpG sites within the CpG groups overlapping the region. This is because nanopolish calls the methylated frequency for each CpG group, not for each CpG site. Since DNA methylation levels in tumor samples are overall lower than normal tissue samples, methylation levels between normal tissue and tumor samples at cancer-specific TE fusion positions cannot be directly compared. Thus, we compared methylation levels of 88 cancer-specific TE fusions from five cancer samples with methylation levels of 1, 295 normal TE fusions using the two-sided Wilcoxon rank sum test.

### Characterization of cancer-specific TE fusions

#### Enrichment analysis of cancer-specific TE fusions in known cancer genes

We obtained the information on tumor suppressor genes and oncogenes from the COSMIC Cancer Gene Census (https://cancer.sanger.ac.uk/census, accessed Sep 16, 2019). We calculated the odds ratio of the cancer gene proportion between normal TE fusions and cancer-specific TE fusions using logistic regression.

#### Survival analysis

We used patient survival data of 201 CoPM cases, 334 ICGC cases, and 9,530 TCGA cases to investigate the association between cancer-specific TE fusions and patient outcome. We calculated overall survival (OS) from the date of diagnosis to the date of death. We selected cancer-specific TE fusions in known cancer genes detected in more than three cancer samples of the same cancer type and calculated the hazard ratio using the Cox proportional hazard model.

#### Immune cell marker expression

We measured gene expression levels (FPKM) in 224 CoPM colon cancer samples, 352 ICGC cancer samples, and 9,619 TCGA cancer samples after upper quantile normalization. We obtained immune cell marker gene sets from a previous study^44^ and measured immune cell activity scores using the ssGSEA^57^ method in the GSVA package^58^. We calculated the Spearman rank correlation between immune cell scores and the corrected number of cancer-specific TE fusions per sample for each cancer type and TE class.

### T-cell immunity analysis of TE-fusion peptides

#### Prediction of MHC-I-binding TE-fusion peptides

Among the cancer-specific TE fusions detected in the CoPM cohort, we searched for events that can induce T-cell immune response by creating neoantigens. First, we selected exonic/exonization type TE fusions as they can alter protein-coding sequences. The coding-sequence change was described as the insertion of the TE sequence into the 5′ breakpoint site to create a variant call format (VCF) file as the input for MuPeXI (ver. 1.2.0) ^41^. We used only the proximal portion of the TE sequence which can be sequenced in short-read RNA sequencing. We used OptiType (ver. 1.3.1) ^59^ and RNA sequencing data to identify HLA genotypes. We used MuPeXI to identify all potential novel peptides, ranging in length from 8 to 11 amino acids, that are not typically found in humans. The likelihood of the binding of the peptide to MHC-I was predicted with netMHCpan (ver. 4.0) ^60^.

#### Structure-based filtering of MHC-I binding TE-fusion peptides

Given the MHC-I and peptide sequences, we selected the most similar template complex structure in the PDB^61^. Our selection was firstly based on the MHC sequence identity and secondly on peptide identity. To match the query sequence, we redesigned the template structure using the “BuildModel” command of FoldX (ver. 5) ^62^. We then relaxed and minimized the backbone structure using the AMBER99sb^63^ with the GB/SA implicit solvent model^64^ using the TINKER molecular dynamics package (ver. 6.3.3) ^65^. The sidechains of the backbone relaxed complex were then re-optimized against the FoldX energy until no energy improvement was made. The “AnalyseComplex’’ command was used to calculate the peptide-MHC-I (pMHC-I) interaction. The calculated interaction energy was normalized for the pre-calculated interaction energy distribution of the target HLA in IEDB^66^. We used the Z-score of the FoldX interaction energy as the pMHC-I binding score (The lower the stronger). Considering the TCR recognition of the bound peptide, we employed FreeSASA (ver. 2.0.4) ^67^ to calculate the solvent accessibility of the bound peptide. If no mutations were solvent-accessible (*i.e.*, not TCR-recognizable), the pMHC was discarded.

#### In vitro assessment of peptide immunogenicity

We evaluated the peptide binding using NeoScreen MHC/Peptide Binding Assays (Immunitrack ApS, Copenhagen, Denmark). The fluorescent peptide–dextramers were synthesized by Immudex, Copenhagen, Denmark. Peripheral blood mononuclear cells (PBMCs) were isolated from blood samples of healthy donors, and their HLA types were analyzed by the hematopoietic stem cell bank at the Catholic University, Seoul, Korea. In brief, PBMCs were cultured in AIM-V medium (Gibco, Thermo Fisher Scientific, Waltham, MA, USA) supplemented with DNase I (Roche, Basel, Switzerland) overnight. The next day, cells were harvested and pulsed with 10 μM peptides. Mitomycin C-treated, peptide-pulsed T-cells were cocultured with PBMCs in the presence of IL-2 (15 U/ml) and IL-15 (5 ng/ml), and cytokines were replenished every week. On day 21, peptide-specific T-cells were stained with pMHC-I dextramer labeled with fluorescein isothiocyanate (Immudex, Copenhagen, Denmark) for 30 min at room temperature in the dark and analyzed by flow cytometry (FACSCanto II, BD, Franklin Lakes, NJ, USA). The experiment involving healthy donor PBMC was conducted in compliance with the ethical regulations and guidelines and was approved by the institutional review board at Catholic University under the IRB number MIRB-00092_3-003.

## Supporting information

Supplementary table STs

## Data availability

The raw RNA-seq and WGS data from the CoPM cohort have been deposited in the Korean Nucleotide Archive (KoNA) at https://kobic.re.kr/kona. The datasets are identified by the following accession IDs: KAP230605, KAP230606, KAP230607, KAP230608, KAP230609, KAP230610, KAP230611, and KAP230612. The long-read ONT WGS FASTQ files for five pairs of colorectal cancer and normal samples, as well as RNA-seq FASTQ files for two cancer cell lines (NCI-H1299 and HCC827), have been deposited in KoNA under the accession ID KAP230581. The methylation data have been deposited in Korea BioData Station (K-BDS, https://kbds.re.kr/) with the project IDs PRJKA2086323, PRJKA2086324, PRJKA2086325, and PRJKA2086326. The merged set of rTea fusion calls from GTEx, TCGA, ICGC, and CoPM samples along with non-redundant TE fusions, is available on https://gitlab.aleelab.net/junseokpark/rtea-results.

## Code availability

The code of *rTea* is available at https://github.com/ealeelab/rtea.

## Acknowledgements

This study was supported by the Suh Kyungbae Foundation (E.A.L) and DP2 AG072437 (E.A.L). This study was additionally supported by the Bio and Medical Technology Development Program of the Korean National Research Foundation funded by the Ministry of Science and ICT (NRF-2017M3A9A7050803, NRF-2020M3A9G3080281, NRF-2020R1A5A2031185, NRF-2021R1F1A103769, NRF-2021M3A9G4078330, and NRF-2021M3A9G407833023).

## Author contributions

E.A.L. conceived and designed the study. B.L. and J.P. developed *rTea*. J.P. optimized *rTea* and established a scalable pipeline on Google Cloud Platform. B.L. and J.P. analyzed TCGA/ICGC and GTEx RNA-seq profiles. B.L. and J.P. characterized TE fusions with statistical advice from Eduardo Maury. B.L. performed splicing, cancer gene enrichment, prognosis, and DNA methylation analysis. Y.K. and J.K.C. contributed to ONT WGS-based DNA methylation analysis. Y.J.K. performed RT-PCR and Sanger sequencing validation. B.L. and Y.C. performed immunogenicity analysis. O.K.H, H.W.L and K.H contributed to in vitro immunogenicity experiments. J.-Y.L., H.-R.S., H.-J.K., J.-W.L., M.-H.J., S.-C.K., H.B.C., D.-W.K., M.K., E.-J.C, and Y.K. generated Illumina and ONT WGS, RNA-seq, and DNA methylation array data for the CoPM consortium. B.L., J.P., Y. C., A.V., and E.A.L wrote the manuscript. E.A.L. and W.-Y.P. supervised the study. All authors reviewed the manuscript.

## References

1. Stewart, C., et al. A comprehensive map of mobile element insertion polymorphisms in humans. PLoS Genet 7, e1002236 (2011).

2. Sudmant, P.H., et al. An integrated map of structural variation in 2,504 human genomes. Nature 526, 75–81 (2015).

3. Borges-Monroy, R., et al. Whole-genome analysis reveals the contribution of non-coding de novo transposon insertions to autism spectrum disorder. Mob DNA 12, 28 (2021).

4. Lee, E., et al. Landscape of somatic retrotransposition in human cancers. Science 337, 967–971 (2012).

5. Tubio, J.M.C., et al. Mobile DNA in cancer. Extensive transduction of nonrepetitive DNA mediated by L1 retrotransposition in cancer genomes. Science 345, 1251343 (2014).

6. Rodriguez-Martin, B., et al. Pan-cancer analysis of whole genomes identifies driver rearrangements promoted by LINE-1 retrotransposition. Nat Genet 52, 306–319 (2020).

7. Chu, C., et al. Comprehensive identification of transposable element insertions using multiple sequencing technologies. Nat Commun 12, 3836 (2021).

8. Scott, E.C., et al. A hot L1 retrotransposon evades somatic repression and initiates human colorectal cancer. Genome Res 26, 745–755 (2016).

9. Cajuso, T., et al. Retrotransposon insertions can initiate colorectal cancer and are associated with poor survival. Nat Commun 10, 4022 (2019).

10. Jang, H.S., et al. Transposable elements drive widespread expression of oncogenes in human cancers. Nature genetics 51, 611–617 (2019).

11. Lock, F.E., et al. A novel isoform of IL-33 revealed by screening for transposable element promoted genes in human colorectal cancer. PLoS One 12, e0180659 (2017).

12. Babaian, A. & Mager, D.L. Endogenous retroviral promoter exaptation in human cancer. Mob DNA 7, 24 (2016).

13. Hancks, D.C., Ewing, A.D., Chen, J.E., Tokunaga, K. & Kazazian, H.H., Jr. Exon-trapping mediated by the human retrotransposon SVA. Genome Res 19, 1983–1991 (2009).

14. Lev-Maor, G., Sorek, R., Shomron, N. & Ast, G. The birth of an alternatively spliced exon: 3’ splice-site selection in Alu exons. Science 300, 1288–1291 (2003).

15. Attig, J., et al. LTR retroelement expansion of the human cancer transcriptome and immunopeptidome revealed by de novo transcript assembly. Genome Res 29, 1578–1590 (2019).

16. Soygur, B. & Sati, L. The role of syncytins in human reproduction and reproductive organ cancers. Reproduction 152, R167–R178 (2016).

17. Reid Cahn, A., Bhardwaj, N. & Vabret, N. Dark genome, bright ideas: Recent approaches to harness transposable elements in immunotherapies. Cancer Cell 40, 792–797 (2022).

18. Kong, Y., et al. Transposable element expression in tumors is associated with immune infiltration and increased antigenicity. Nat Commun 10, 5228 (2019).

19. Smith, C.C., et al. Endogenous retroviral signatures predict immunotherapy response in clear cell renal cell carcinoma. J Clin Invest 128, 4804–4820 (2018).

20. Shah, N.M., et al. Pan-cancer analysis identifies tumor-specific antigens derived from transposable elements. Nat Genet 55, 631–639 (2023).

21. Merlotti, A., et al. Noncanonical splicing junctions between exons and transposable elements represent a source of immunogenic recurrent neo-antigens in patients with lung cancer. Sci Immunol 8, eabm6359 (2023).

22. Ewing, A.D. Transposable element detection from whole genome sequence data. Mob DNA 6, 24 (2015).

23. Barretina, J., et al. The Cancer Cell Line Encyclopedia enables predictive modelling of anticancer drug sensitivity. Nature 483, 603–607 (2012).

24. The Cancer Genome Atlas Research Network, et al. The Cancer Genome Atlas Pan-Cancer analysis project. Nat Genet 45, 1113-1120 (2013).

25. The ICGC/TCGA Pan-Cancer Analysis of Whole Genomes Consortium. Pan-cancer analysis of whole genomes. Nature 578, 82-93 (2020).

26. Lonsdale, J., et al. The genotype-tissue expression (GTEx) project. Nature genetics 45, 580–585 (2013).

27. GTEx Consortium. Human genomics. The Genotype-Tissue Expression (GTEx) pilot analysis: multitissue gene regulation in humans. Science 348, 648–660 (2015).

28. Mele, M., et al. Human genomics. The human transcriptome across tissues and individuals. Science 348, 660–665 (2015).

29. Simpson, A.J., Caballero, O.L., Jungbluth, A., Chen, Y.T. & Old, L.J. Cancer/testis antigens, gametogenesis and cancer. Nat Rev Cancer 5, 615–625 (2005).

30. Gorbunova, V., et al. The role of retrotransposable elements in ageing and age-associated diseases. Nature 596, 43–53 (2021).

31. Spinelli, R., et al. Molecular basis of ageing in chronic metabolic diseases. J Endocrinol Invest 43, 1373–1389 (2020).

32. Gardner, E.J., et al. The Mobile Element Locator Tool (MELT): population-scale mobile element discovery and biology. Genome Res 27, 1916–1929 (2017).

33. Jung, H., Choi, J.K. & Lee, E.A. Immune signatures correlate with L1 retrotransposition in gastrointestinal cancers. Genome Res 28, 1136–1146 (2018).

34. Burns, K.H. Transposable elements in cancer. Nat Rev Cancer 17, 415–424 (2017).

35. Miki, Y., et al. Disruption of the APC gene by a retrotransposal insertion of L1 sequence in a colon cancer. Cancer Res 52, 643–645 (1992).

36. Sorek, R., et al. Minimal conditions for exonization of intronic sequences: 5’ splice site formation in alu exons. Mol Cell 14, 221–231 (2004).

37. Conti, A., et al. Identification of RNA polymerase III-transcribed Alu loci by computational screening of RNA-Seq data. Nucleic Acids Res 43, 817–835 (2015).

38. Hur, K., et al. Hypomethylation of long interspersed nuclear element-1 (LINE-1) leads to activation of proto-oncogenes in human colorectal cancer metastasis. Gut 63, 635–646 (2014).

39. Miglio, U., et al. The expression of LINE1-MET chimeric transcript identifies a subgroup of aggressive breast cancers. Int J Cancer 143, 2838–2848 (2018).

40. Rashed, W.M., et al. MET canonical transcript expression is a predictive biomarker for chemo-sensitivity to MET-inhibitors in hepatocellular carcinoma cell lines. J Cancer Res Clin Oncol 147, 167–175 (2021).

41. Bjerregaard, A.M., Nielsen, M., Hadrup, S.R., Szallasi, Z. & Eklund, A.C. MuPeXI: prediction of neo-epitopes from tumor sequencing data. Cancer Immunol Immunother 66, 1123–1130 (2017).

42. Chen, R., Ishak, C.A. & De Carvalho, D.D. Endogenous Retroelements and the Viral Mimicry Response in Cancer Therapy and Cellular Homeostasis. Cancer Discov 11, 2707–2725 (2021).

43. Barbie, D.A., et al. Systematic RNA interference reveals that oncogenic KRAS-driven cancers require TBK1. Nature 462, 108–112 (2009).

44. Rooney, M.S., Shukla, S.A., Wu, C.J., Getz, G. & Hacohen, N. Molecular and genetic properties of tumors associated with local immune cytolytic activity. Cell 160, 48–61 (2015).

45. de Cubas, A.A., et al. DNA hypomethylation promotes transposable element expression and activation of immune signaling in renal cell cancer. JCI Insight 5(2020).

46. Mehdipour, P., et al. Epigenetic therapy induces transcription of inverted SINEs and ADAR1 dependency. Nature 588, 169–173 (2020).

47. Roulois, D., et al. DNA-Demethylating Agents Target Colorectal Cancer Cells by Inducing Viral Mimicry by Endogenous Transcripts. Cell 162, 961–973 (2015).

48. Smith, C.C., et al. Endogenous retroviral signatures predict immunotherapy response in clear cell renal cell carcinoma. The Journal of clinical investigation 128, 4804–4820 (2019).

49. Modzelewski, A.J., et al. A mouse-specific retrotransposon drives a conserved Cdk2ap1 isoform essential for development. Cell 184, 5541–5558 e5522 (2021).

50. Chen, S., Zhou, Y., Chen, Y. & Gu, J. fastp: an ultra-fast all-in-one FASTQ preprocessor. Bioinformatics 34, i884–i890 (2018).

51. Kim, D., Paggi, J.M., Park, C., Bennett, C. & Salzberg, S.L. Graph-based genome alignment and genotyping with HISAT2 and HISAT-genotype. Nat Biotechnol 37, 907–915 (2019).

52. Li, H., et al. The Sequence Alignment/Map format and SAMtools. Bioinformatics 25, 2078–2079 (2009).

53. Li, H. & Durbin, R. Fast and accurate short read alignment with Burrows-Wheeler transform. Bioinformatics 25, 1754–1760 (2009).

54. Bao, W., Kojima, K.K. & Kohany, O. Repbase Update, a database of repetitive elements in eukaryotic genomes. Mob DNA 6, 11 (2015).

55. Fernandes, J.D., et al. The UCSC repeat browser allows discovery and visualization of evolutionary conflict across repeat families. Mob DNA 11, 13 (2020).

56. Huang, X. & Madan, A. CAP3: A DNA sequence assembly program. Genome Res 9, 868–877 (1999).

57. Barbie, D.A., et al. Systematic RNA interference reveals that oncogenic KRAS-driven cancers require TBK1. Nature 462, 108–112 (2009).

58. Hanzelmann, S., Castelo, R. & Guinney, J. GSVA: gene set variation analysis for microarray and RNA-seq data. BMC Bioinformatics 14, 7 (2013).

59. Szolek, A., et al. OptiType: precision HLA typing from next-generation sequencing data. Bioinformatics 30, 3310–3316 (2014).

60. Jurtz, V., et al. NetMHCpan-4.0: Improved Peptide-MHC Class I Interaction Predictions Integrating Eluted Ligand and Peptide Binding Affinity Data. J Immunol 199, 3360–3368 (2017).

61. Berman, H.M., et al. The Protein Data Bank. Nucleic Acids Res 28, 235–242 (2000).

62. Schymkowitz, J., et al. The FoldX web server: an online force field. Nucleic Acids Res 33, W382–388 (2005).

63. Lindorff-Larsen, K., et al. Improved side-chain torsion potentials for the Amber ff99SB protein force field. Proteins 78, 1950–1958 (2010).

64. Onufriev, A., Bashford, D. & Case, D.A. Modification of the generalized Born model suitable for macromolecules. The Journal of Physical Chemistry B 104, 3712–3720 (2000).

65. Rackers, J.A., et al. Tinker 8: Software Tools for Molecular Design. J Chem Theory Comput 14, 5273–5289 (2018).

66. Vita, R., et al. The Immune Epitope Database (IEDB): 2018 update. Nucleic Acids Res 47, D339–D343 (2019).

67. Mitternacht, S. FreeSASA: An open source C library for solvent accessible surface area calculations. F1000Res 5, 189 (2016).

