## Supplementary figures and tables for "Pan-cancer analysis reveals multifaceted roles of retrotransposon-fusion RNAs"

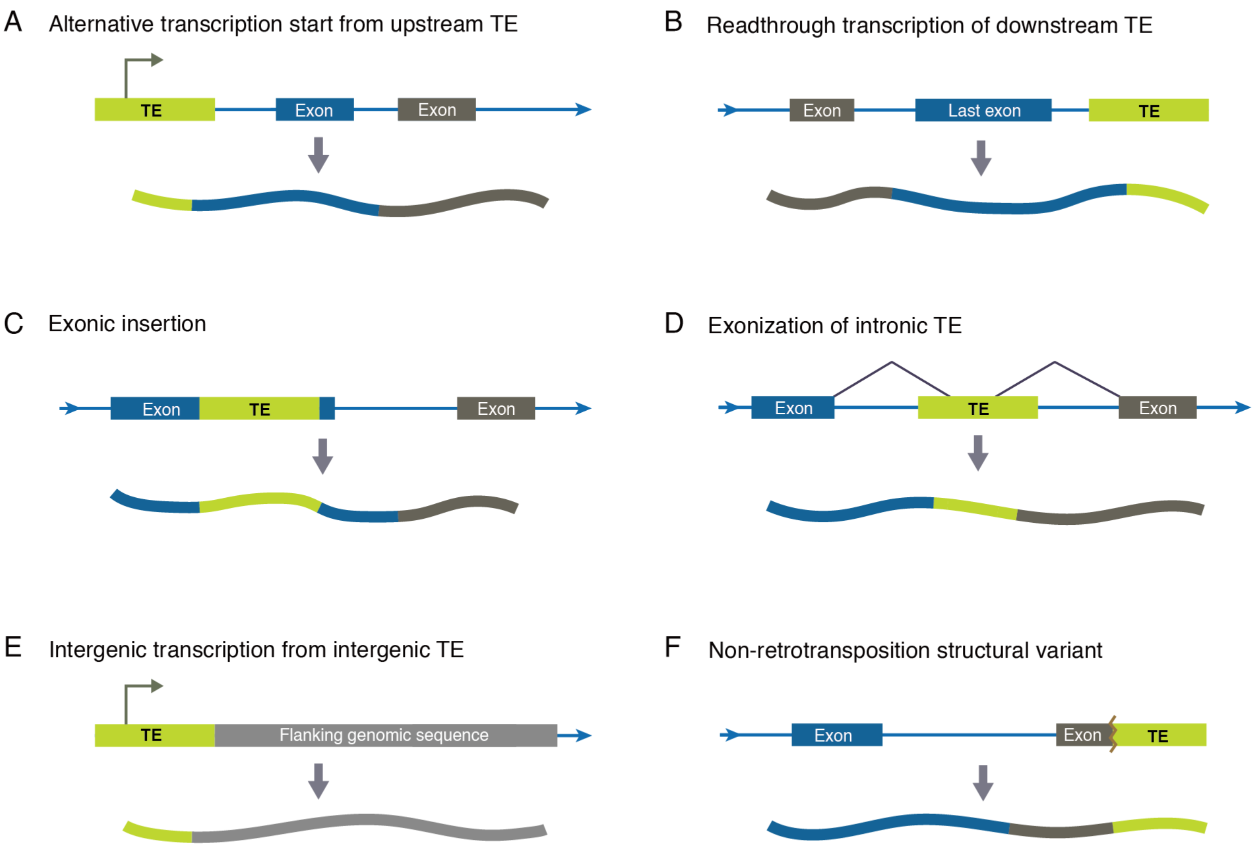


**Figure S1. Mechanisms underlying different types of TE fusion transcripts.** (A) Transcription begins at promoters within the TE body and continues into the gene sequence by readthrough or splicing. (B) Transcription begins within the gene and continues into a downstream TE via readthrough. (C) The TE inserted within an exon is co-expressed with genes (D) An intronic TE is spliced into the mature transcript through exonization. (E) An intergenic TE expresses along with the downstream genomic sequence. (F) Chromosomal rearrangement unrelated to retrotransposition forms a fusion between the gene and TE.

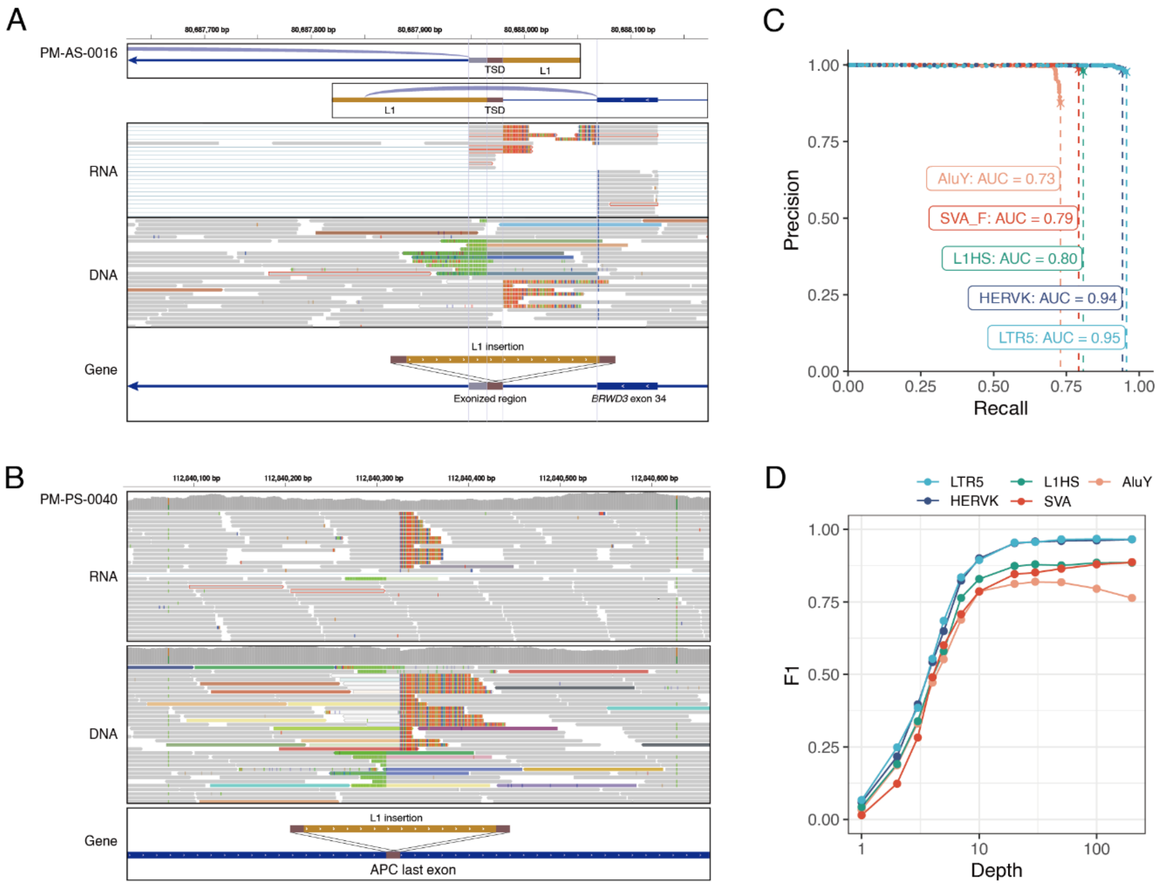


**Figure S2. Performance evaluation of *rTea* by *in silico* simulation.** (A) An example TE fusion detected by *rTea* involving the exonization of a non-reference L1 inserted into an intron from the sample (PM-AS-0016-T-A1) of CoPM. The top track shows the RNA coverage on the reference genome around the L1 insertion, including a portion of the exonized reference sequence downstream of the insertion. The magnified region shows the RNA reads supporting the exonization and the DNA reads supporting the L1 insertion. (B) An example TE fusion detected by *rTea* involving a non-reference L1 inserted within an exon from the sample (PM-PS-0040-T-A2) of CoPM. The TE fusion is seen in both the DNA and RNA sequencing data. (C) Precision-recall curve and area under the curve (AUC) for the *in silico* simulation of *rTea* by TE family. The AUC was obtained by varying the cutoff from 1X to 200X with read length 100bp (167 as insert size) data. (D) The F1 values for *rTea* results by depth of sequencing per TE family based on *in silico* simulation.

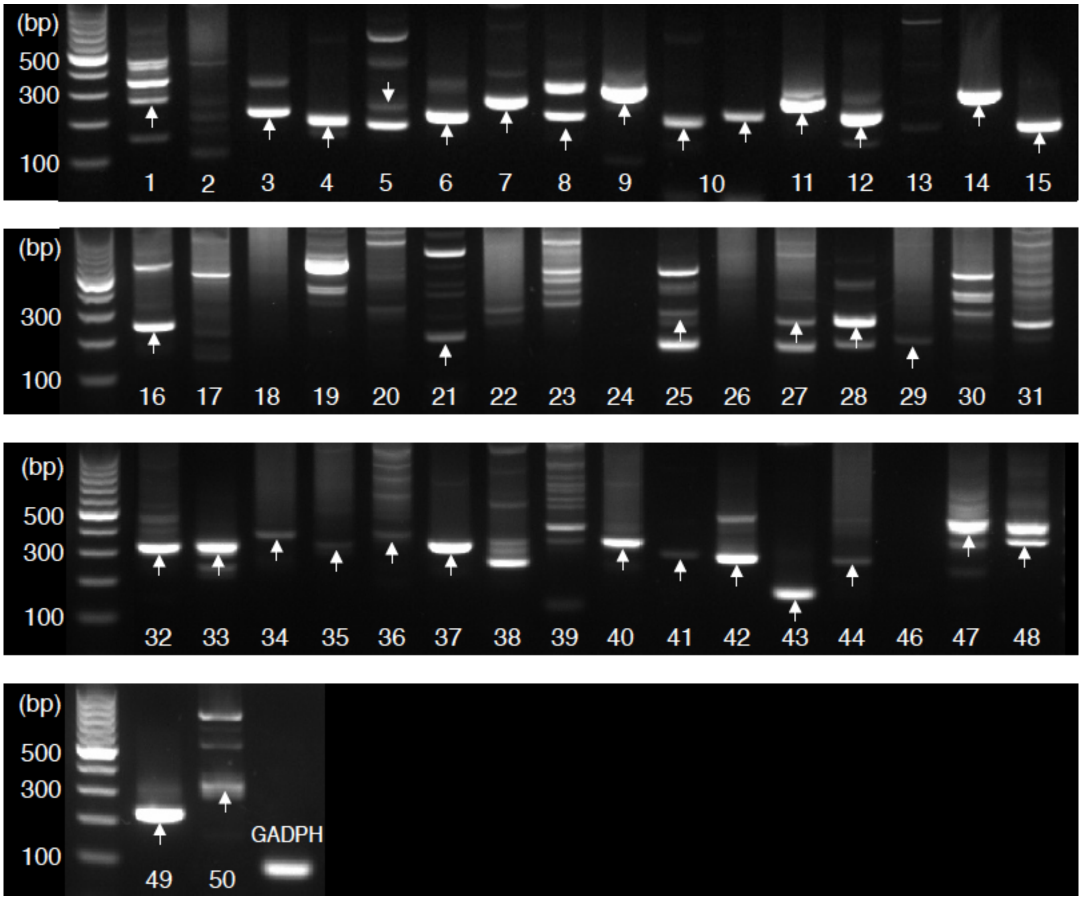


**Figure S3. RT-PCR validation of *rTea* results using the H1299 cell line.** We produced RNA-seq data using the H1299 cell line (breast cancer) and performed RT-PCR for 50 randomly selected TE fusions detected by *rTea*. **Table ST2** describes the details of each TE fusion selected for validation. A white arrow indicates a band at the expected fragment size for the fusion. Among the 50 TE fusions, 24 (48%) showed clear bands, 10 (20%) showed multiple bands including the expected band, 15 (30%) showed no band, and PCR failed for 1 (2%) TE fusion.

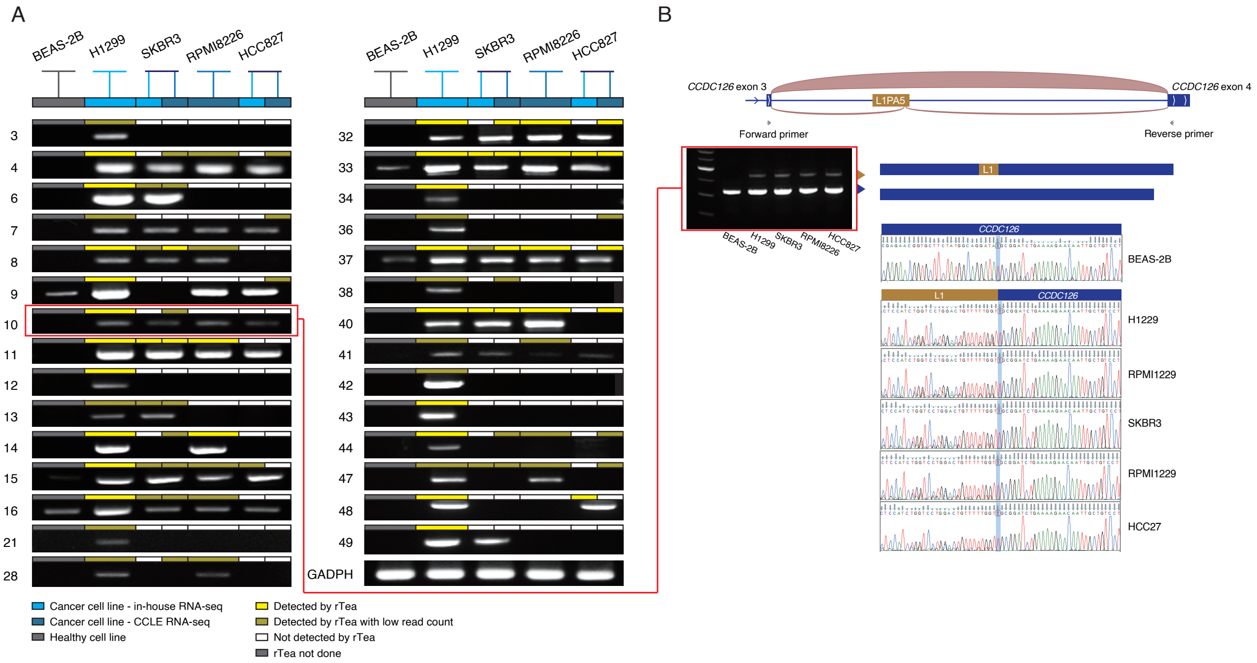


**Figure S4. RT-PCR validation of TE fusions across 5 cell lines.** (A) To eliminate the possibility of non-specific PCR products, we conducted RT-PCR on the TE fusions validated in H1299 with four additional cell lines (SKBR3, breast cancer; RPMI8226, HCC827, multiple myeloma; and BEAS-2B, normal bronchial epithelium). We ran *rTea* on all cell lines except BEAS-2B using a combination of direct RNA-sequencing data and data from CCLE (Cancer Cell Line Encyclopedia). Among the 7 TE fusions detected by *rTea* only in H1299 (3, 12, 21, 34, 42, 43, 49), 6 TE fusions showed PCR product only in H1299. (B) RT-PCR and Sanger sequencing verification of the exonization of an intronic L1 within *CCDC126* detected by *rTea*. Among the PCR bands, the orange arrow indicates L1PA5 exonization and the blue arrow corresponds to the typical *CCDC126* transcript. Sanger sequencing confirmed the L1 exonization sequence.

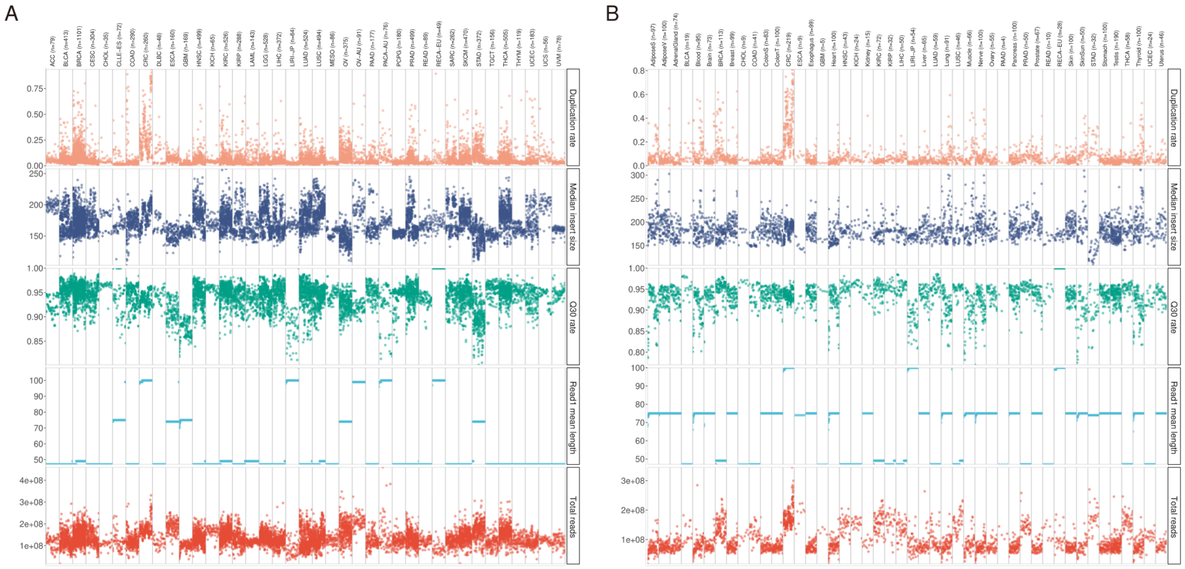

**Figure S5. Highly variable technical factors that affect TE fusion detection.** PCR duplication rate, median insertion size, Q30 rate, read length, and total read count are shown for each sample grouped by cancer or tissue type from (A) TCGA, ICGC, and CoPM, and (B) GTEx data.

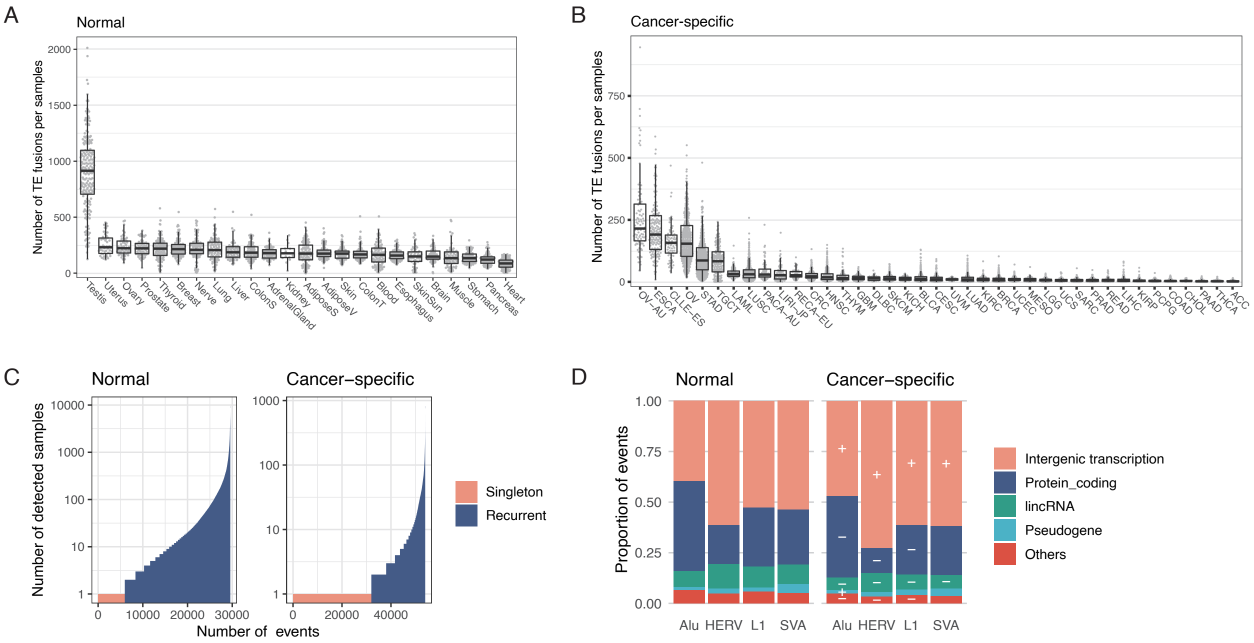


**Figure S6. Characterization of TE fusions.** (A) The number of TE fusions per sample in GTEx data before correcting the means for technical variables. (B) The number of cancer-specific TE fusions per sample before correction of technical variables. (C) Distribution of the number of samples each TE fusion was detected in, with TEs detected only in a single sample (singleton, orange) and detected in > 2 samples (recurrent, blue). (D) The proportion of partner gene types involved in TE fusions. The gene types significantly more common in cancer-specific TE fusions were marked by '+' and gene types with significantly less common in cancer-specific TE fusions were marked by '-' based on Fisher's exact test with FDR correction.

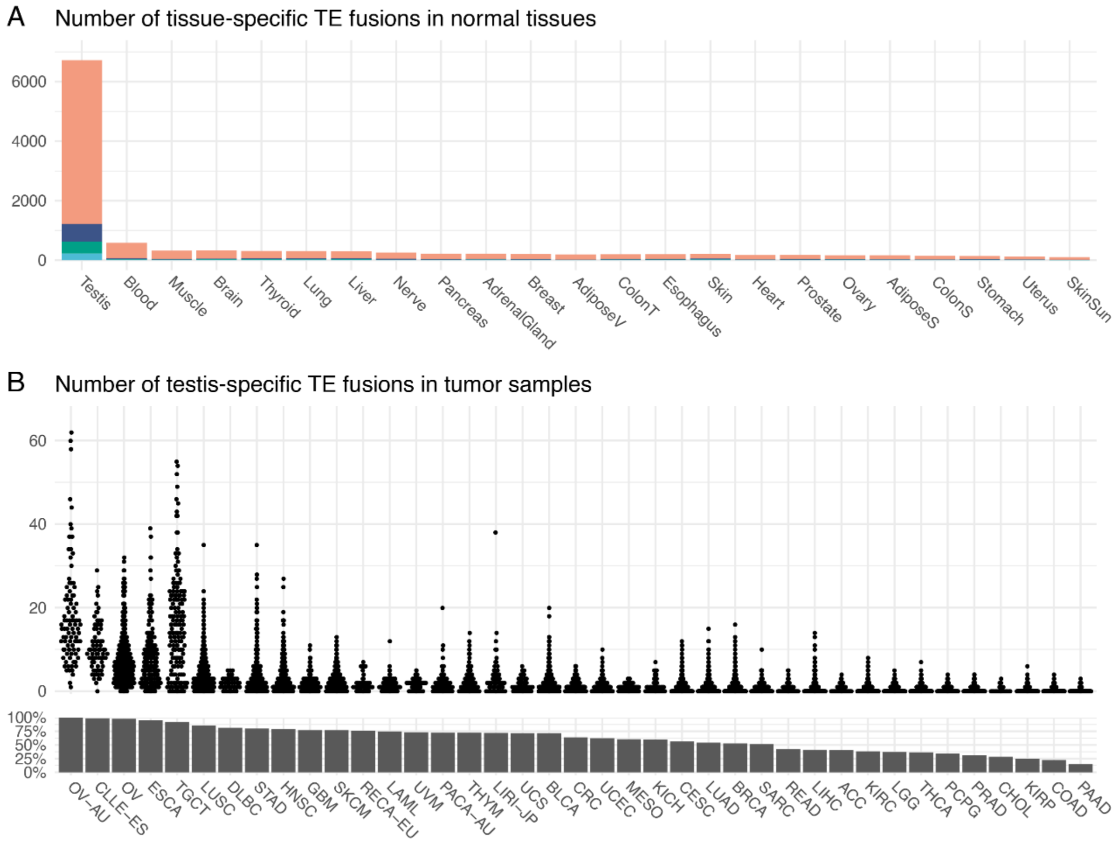


**Figure S7. Abundant testis-specific TE fusions.** (A) TE fusions detected only in specific tissues are counted. Testis had considerably large number of tissue-specific TE fusions. (B) Testis-specific TE fusions were detected in various types of cancers. Number of detected testis-specific TE fusions per sample are displayed for each cancer type. Bar graph on the lower pane shows the percent of samples expressing testis-specific TE fusions for each cancer type.

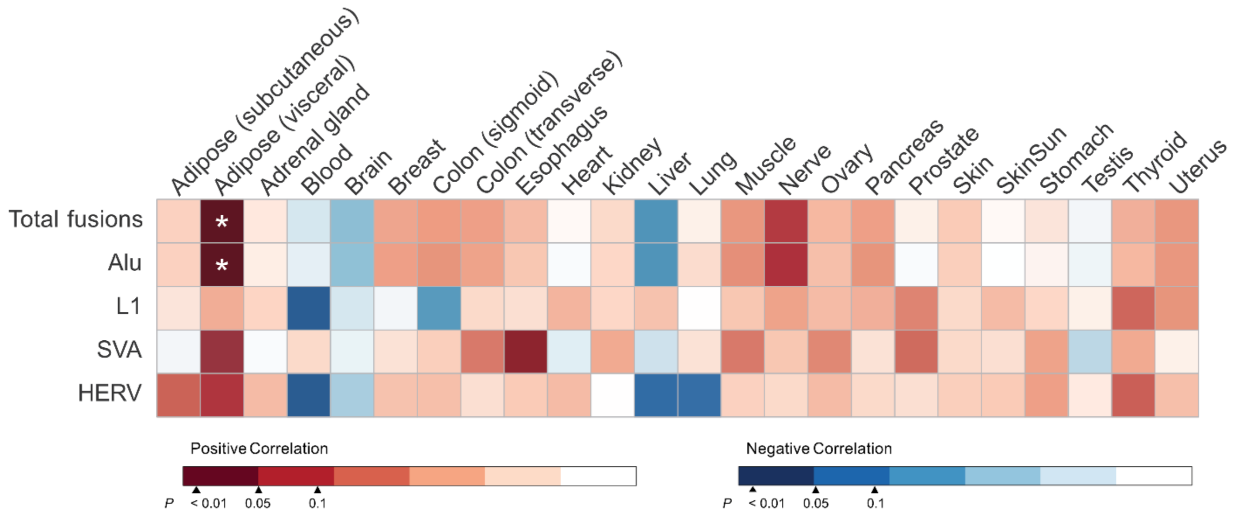


**Figure S8. Association between TE fusion counts and donor ages.** Significant correlation was marked with '*' (FDR < 0.05, negative binomial mixed models).

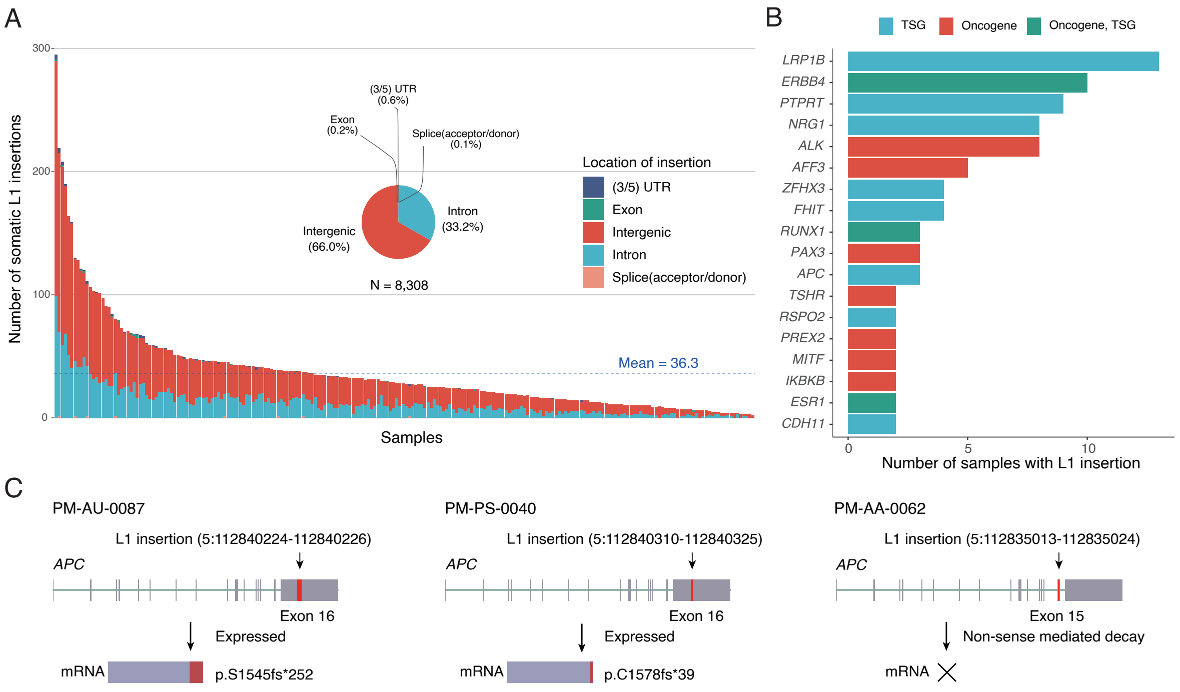


**Figure S9. Somatic L1 insertions in cancers genes, including *APC*, in CoPM colorectal cancer samples.** (A) Number of somatic L1 detected in WGS data from 229 CoPM colorectal cancer samples. The bars represent the union of xTEA, MELT, and TraFiC insertion calls, color coded by genomic annotation. The pie chart shows the percentage of insertions for each genomic category. (B) Recurrent somatic L1 insertions in known cancer genes. The number of cancer patients with at least one somatic L1 insertion is depicted for each gene from oncogenes, tumor suppressor genes, and both categories annotated in the COSMIC Cancer Gene Census by red , blue and green bar, respectively. (C) Somatic L1 insertions in *APC* in three patients and their RNA expression. Only L1 retrotranspositions in the last exon 16 were detected in RNA-seq data.

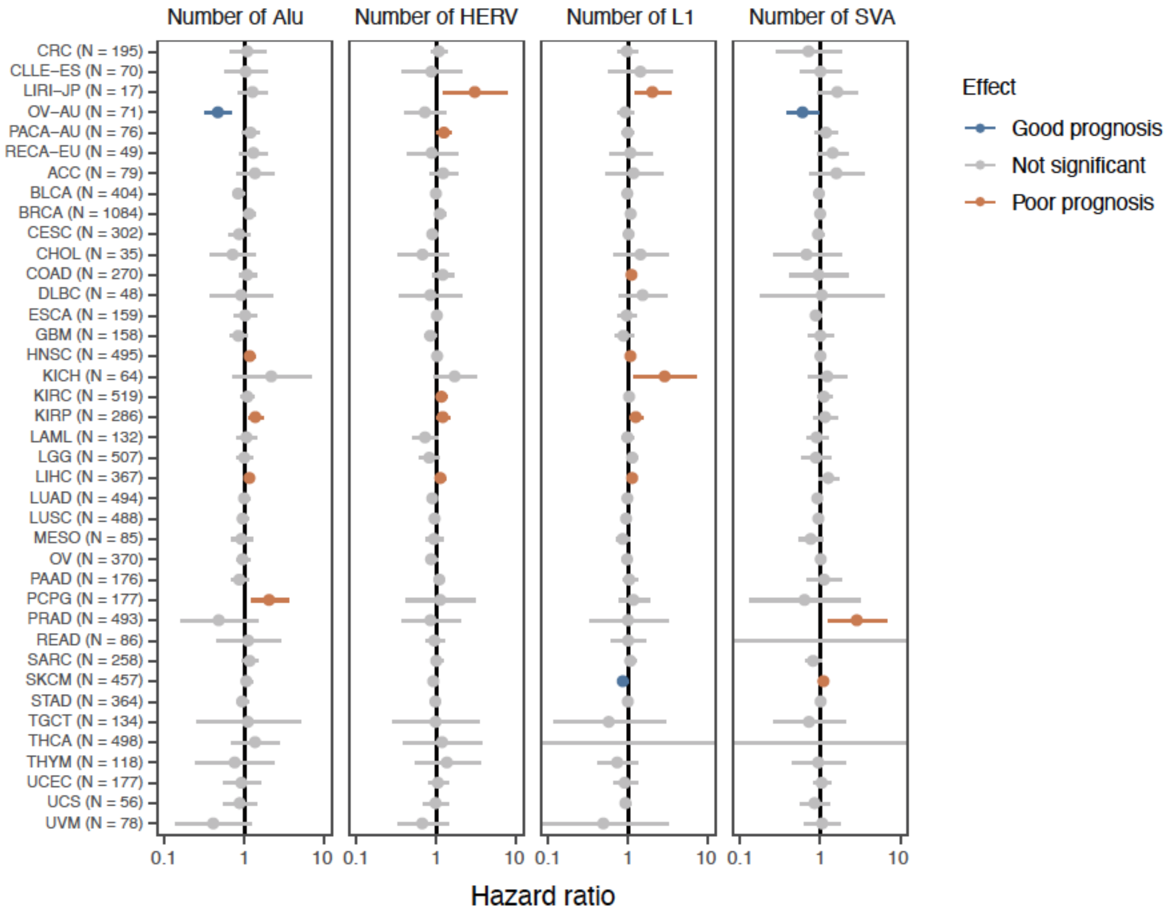


**Figure S10. Survival outcome according to the number of cancer-specific TE fusions.** The hazard ratio was calculated using the Cox proportional hazard model with number of recurrent (present in ≥ 3 samples) cancer-specific TE fusions in cancer related genes per sample for each cancer type in TCGA and ICGC and patient survival data. Colored ranges indicate significantly worse (orange) or better (blue) prognosis.

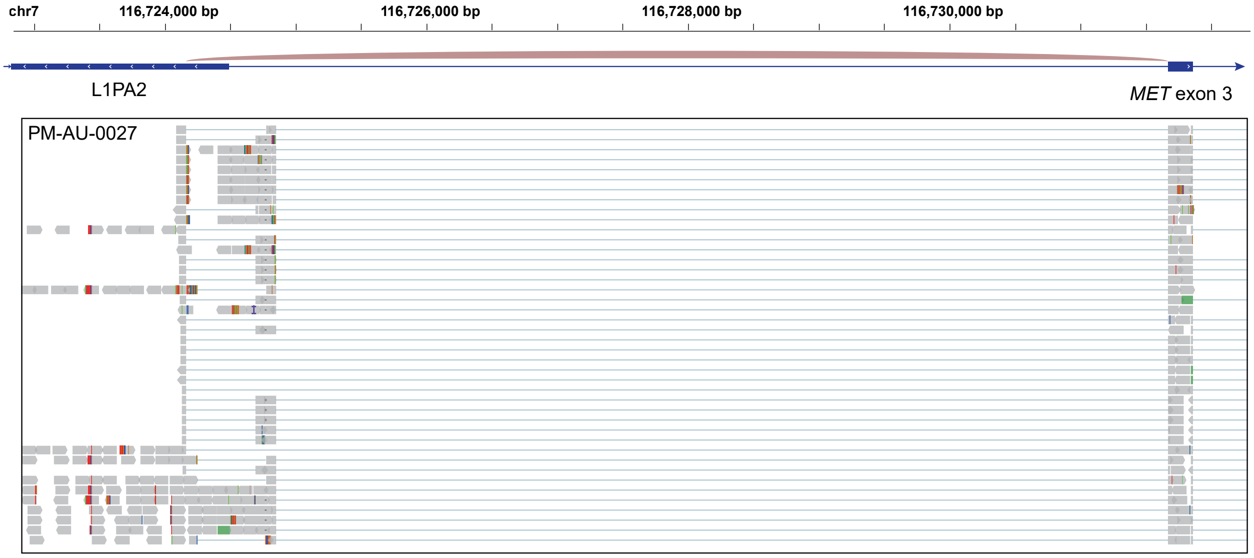


**Figure S11. Genome browser visualization of an alternative transcription start site located within an L1PA2 in *MET* intron 2.** Alternative transcription starting from this L1PA2 has been previously reported.

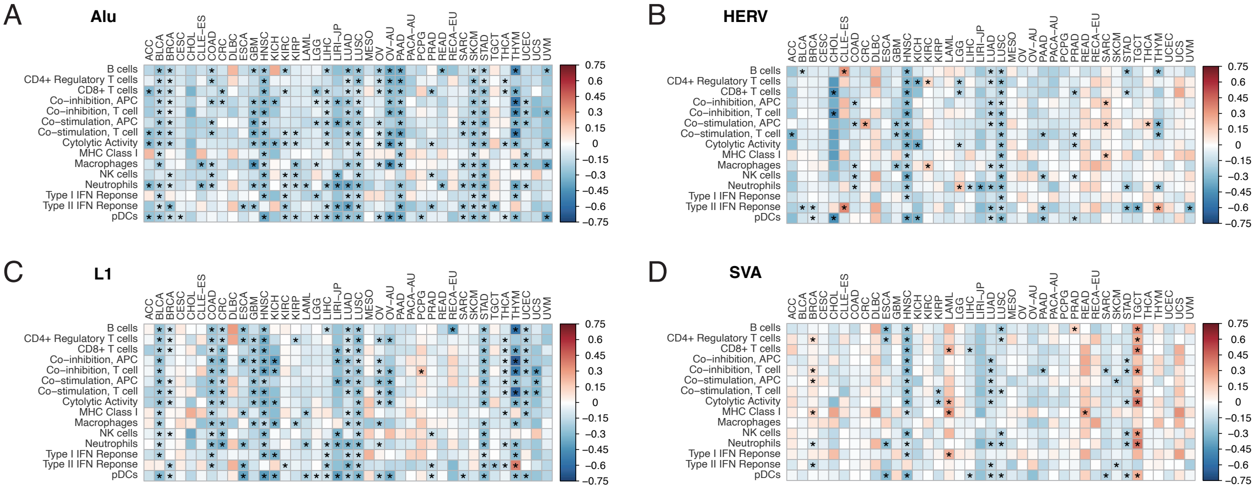


**Figure S12. Correlation between the number of cancer-specific TE fusions and immune score.** Correlation between the number of TE fusions and immune score was shown for (A) Alu, (B) HERV, (C) L1, and (D) SVA. FDR < 0.05 based on Spearman rank correlation marked with a star.

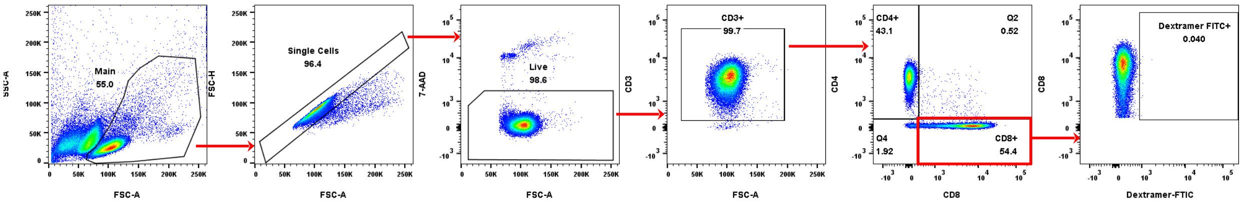


**Figure S13. Gating strategy for flow cytometry analysis.** The gating was performed as follows: (1) The main population of lymphocytes was gated on the dot plot (X axis: FSC-A, Y axis: SSC-A). (2) The single cells were gated using the main population gating set in step 1 on the dot plot (X axis: FSC-A, Y axis: FSC-H). (3) Live cells were gated using the single cell gating set in step 2 on the dot plot (X axis: FSC-A, Y axis: PerCP) (Dead cells are stained with 7-AAD, which is not fluorescent in live cells. Evaluation of 7-AAD staining is possible in the PerCP fluorescence channel). (4) CD3+ cells were gated using the single cell gating set in step 3 on the dot plot (X axis: FSC-A, Y axis: CD3-PE-Cy7). (5) The CD4+ CD8+ T cell region was gated as a quadrant using the CD3+ gating set in step 4 on the dot plot (X axis: CD8-APC, Y axis: CD4-BV421). (6) The Dextramer+ region was gated using the CD8+ T cell region set in step 5 on the dot plot (X axis: Dextramer-FITC, Y axis: CD8-APC).

### Supplementary Tables

**Table S1. Comparison of TE sources between normal TE fusions and cancer-specific TE fusions.**

| Class | Source of TE | Normal TE fusions Count (percent) | Cancer-Specific TE fusions Count (percent) | OR | P | FDR |
|---|---|---|---|---|---|---|
| Alu | Reference genome | 5,070 (80%) | 1,194 (60%) | 0.378 | 1.21.E-71 | 2.67.E-70 |
| | Germline TE | 399 (6%) | 123 (6%) | 0.981 | 8.97.E-01 | 1.00.E+00 |
| | Germline SV | 122 (2%) | 35 (2%) | 0.912 | 7.04.E-01 | 1.00.E+00 |
| | Somatic TEI | 0 (0%) | 1 (0%) | Inf | 5.40.E-01 | 1.00.E+00 |
| | Somatic SV | 0 (0%) | 38 (2%) | Inf | 2.31.E-27 | 5.09.E-26 |
| | Not determined | 767 (12%) | 605 (30%) | 3.170 | 7.76.E-82 | 1.71.E-80 |
| HERV | Reference genome | 327 (78%) | 231 (70%) | 0.685 | 2.98.E-02 | 6.56.E-01 |
| | Germline TE | 8 (2%) | 3 (1%) | 0.477 | 4.20.E-01 | 1.00.E+00 |
| | Germline SV | 15 (4%) | 3 (1%) | 0.250 | 3.51.E-02 | 7.72.E-01 |
| | Somatic SV | 0 (0%) | 5 (2%) | Inf | 3.67.E-02 | 8.06.E-01 |
| | Not determined | 71 (17%) | 86 (26%) | 1.752 | 2.44.E-03 | 5.38.E-02 |
| L1 | Reference genome | 475 (73%) | 434 (70%) | 0.864 | 2.68.E-01 | 1.00.E+00 |
| | Germline TE | 75 (12%) | 37 (6%) | 0.487 | 6.95.E-04 | 1.53.E-02 |
| | Germline SV | 24 (4%) | 6 (1%) | 0.255 | 2.64.E-03 | 5.82.E-02 |
| | Somatic TEI | 0 (0%) | 19 (3%) | Inf | 1.96.E-05 | 4.31.E-04 |
| | Somatic SV | 0 (0%) | 13 (2%) | Inf | 5.92.E-04 | 1.30.E-02 |
| | Not determined | 74 (11%) | 108 (18%) | 1.646 | 2.68.E-03 | 5.90.E-02 |
| SVA | Reference genome | 81 (52%) | 23 (52%) | 1.001 | 1.00.E+00 | 1.00.E+00 |
| | Germline TE | 33 (21%) | 9 (20%) | 0.951 | 1.00.E+00 | 1.00.E+00 |
| | Germline SV | 19 (12%) | 1 (2%) | 0.166 | 9.69.E-02 | 1.00.E+00 |
| | Somatic TEI | 0 (0%) | 1 (2%) | Inf | 5.00.E-01 | 1.00.E+00 |
| | Not determined | 22 (14%) | 10 (23%) | 1.778 | 2.60.E-01 | 1.00.E+00 |

P-values based on Chi-squared tests and corrected for FDR.

**Table S2. Z-score of methylation of the source TE derived from methylation array data.**

| TE class | TE fusion type | Number | Percentage of negative Z-score | Median Z-score | P | FDR |
|---|---|---|---|---|---|---|
| Alu | Intergenic transcription | 2,265 | 49% | 0.038 | 0.139 | 1.000 |
| | Alternative transcription start | 494 | 50% | -0.017 | 0.035 | 0.562 |
| | Exonic/exonization | 2,850 | 48% | 0.042 | 0.332 | 1.000 |
| | Readthrough transcription | 729 | 49% | 0.032 | 0.080 | 1.000 |
| HERV | Intergenic transcription | 224 | 57% | -0.184 | 0.004 | 0.062 |
| | Alternative transcription start | 109 | 67% | -0.340 | 0.004 | 0.059 |
| | Exonic/exonization | 279 | 76% | -0.570 | < 0.001 | < 0.001 |
| | Readthrough transcription | 7 | 14% | 0.650 | 0.297 | 1.000 |
| L1 | Intergenic transcription | 529 | 89% | -1.094 | < 0.001 | < 0.001 |
| | Alternative transcription start | 83 | 78% | -0.591 | < 0.001 | < 0.001 |
| | Exonic/exonization | 348 | 79% | -1.022 | < 0.001 | < 0.001 |
| | Readthrough transcription | 18 | 56% | -0.240 | 0.495 | 1.000 |
| SVA | Intergenic transcription | 113 | 57% | -0.286 | 0.004 | 0.072 |
| | Alternative transcription start | 10 | 60% | -0.440 | 0.375 | 1.000 |
| | Exonic/exonization | 59 | 83% | -2.979 | < 0.001 | < 0.001 |
| | Readthrough transcription | 3 | 33% | 0.416 | 0.750 | 1.000 |

P-values based on Wilcoxon rank sum tests and corrected for FDR.

**Table S3. DNA methylation level of the source TE derived from ONT data.**

| Class | Region | N | Median methylation normal tissue | Median methylation tumor | P |
|---|---|---|---|---|---|
| Alu | Upstream | 45 | 0.84 | 0.85 | 0.769 |
| | TE body | 45 | 0.89 | 0.87 | 0.142 |
| | Downstream | 45 | 0.84 | 0.83 | 0.058 |
| L1 | Upstream | 28 | 0.80 | 0.77 | 0.003 |
| | TE body | 28 | 0.82 | 0.59 | < 0.001 |
| | Downstream | 28 | 0.78 | 0.75 | 0.078 |
| HERV | Upstream | 7 | 0.67 | 0.43 | 0.016 |
| | TE body | 7 | 0.81 | 0.48 | 0.031 |
| | Downstream | 7 | 0.87 | 0.36 | 0.219 |

P-values based on Wilcoxon rank sum tests.
